# Graph-theoretical Analysis of EEG Functional Connectivity during Balance Perturbation in Traumatic Brain Injury: A Pilot Study

**DOI:** 10.1101/2020.10.08.332353

**Authors:** Vikram Shenoy Handiru, Alaleh Alivar, Armand Hoxha, Soha Saleh, Easter S. Suviseshamuthu, Guang Yue, Didier Allexandre

## Abstract

Traumatic Brain Injury (TBI) often results in balance impairment, increasing the risk of falls, and the chances of further injuries. However, the underlying neurophysiological mechanisms of postural control after TBI are not well understood. To this end, we conducted a pilot study with a multimodal approach of EEG, MRI, and Diffusion Tensor Imaging (DTI) to explore the neural mechanisms of unpredictable balance perturbations in 17 chronic TBI participants and 15 matched Healthy Controls (HC). As quantitative measures of the functional integration and segregation of the brain networks during the postural task, we computed the global graph-theoretic network measures (global efficiency and modularity) of brain functional connectivity derived from source-space EEG in different frequency bands. We observed that the TBI group showed a lower balance performance as measured by the Center of Pressure (COP) displacement during the task, and the Berg Balance Scale. They also showed altered brain activation and connectivity during the balance task. In particular, the task modulation of brain network segregation in alpha-band was reduced in TBI. Moreover, the DTI findings revealed that the structural damage is associated with reduced network connectivity and integration. In terms of the neural correlates, we observed a distinct role played by different frequency bands; greater theta-band modularity during the task was strongly correlated with the BBS in TBI group; alpha-band and beta-band graph-theoretic measures were associated with the measures of white matter structural integrity. Our future studies will focus on how postural training will modulate the functional brain networks in TBI.

## Introduction

Traumatic Brain Injury (TBI) is a significant medical and health problem in the United States, with an estimated 2.8 million people sustaining a TBI every year [Taylor et al., 2017]. One of the immediate consequences of TBI is an elevated risk of balance deficit or the loss of postural control [Kaufman et al., 2006] [Pickett, 2007]. Yet, the pathomechanisms of the postural instability remain poorly understood. To develop effective rehabilitation strategies and prognosticating tools for motor function recovery, we must delve into the neurophysiological processes involved in postural control.

One aspect that could guide us in this direction is the “functional connectivity” between the cortical regions involved in postural control during the postural control task. Functional connectivity (FC) refers to the statistical interdependencies between the physiological time series of two regions. In the event of a brain injury, the cortical lesion or the axonal damage can disrupt not only the structural integrity of the white matter (WM) system but also the FC between regions [Gratton et al., 2012]. Regardless of the type of network (i.e., the physical synaptic connection between the set of neurons or the functional connection between activities of two brain regions), a graph theory-based approach is a promising tool to quantify the FC network organization wherein the whole brain is viewed as a graph with several nodes (e.g., anatomically parcellated region). Using these quantifiable measures to identify the cortical biomarkers of balance deficits in TBI could better guide us in developing intervention strategies. This brings us to three research questions:

1. *How do the functional networks pertaining to the balance perturbation task change due to* TBI ?
2. *Are the global task-specific* connectivity *measures derived using graph-theory highlight specific brain functional impairment related to balance deficits in TBI?*
3. *Are task-based brain functional connectivity measures associated with the structural integrity of white matter?*

In pursuit of answers to these questions, we look at the graph-theoretical perspective of brain networks. In this framework, the brain connectivity graph is considered as a network made of nodes and edges. Each node corresponds to a cortical region of interest (based on anatomical/functional parcellation) and the edge corresponds to the strength of the functional connectivity between any two connected nodes. Using these fundamental elements of a graph network, graph theory aims at computing several metrics (i.e., node degree, local and global efficiency, clustering coefficient, modularity, etc.) to characterize and study the functional or structural organization of a network of nodes and edges to better understand the neuroanatomical and neurophysiological basis of brain function. A graphical illustration is presented in Fig. 1 to describe the basic concepts of network neuroscience related to this study.

**Figure 1:**
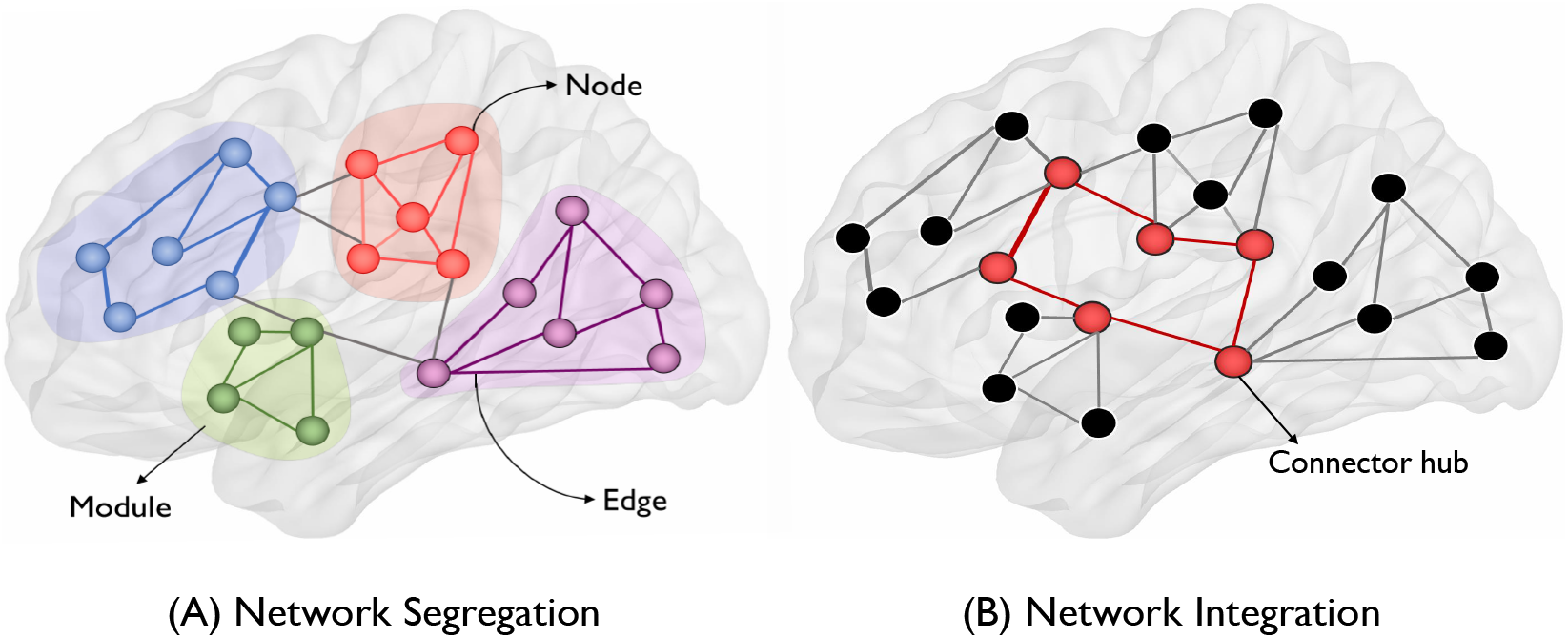
Illustration of graph-theoretic measures. In the context of brain networks in this study, every anatomical region-of-interest is a *node,* whereas the connection between nodes is termed as an *edge.* Colored shaded regions correspond to *modules* which are groups of interconnected nodes but have fewer connections to other modules. On the left, the concept of *network segregation* is presented as a network comprising of multiple modules (or segregated sub-networks) whereas, on the right, the *network integration* is illustrated as a group of distant regions interconnected by long-range connections via connector hubs (nodes with a high degree of connectivity).

In particular, the concept of *functional integration* and *functional segregation* is gaining significant attention in the network neuroscience community [Newman, 2006]; [Stam, 2014]; [Rubinov and Sporns, 2010]; [Hagmann et al., 2008]; [Cohen and D’Esposito, 2016]. In this conceptual framework, as shown in Fig. 1, the connectivity structure of the brain is considered to be organized in several statistically independent clusters of nodes or subnetworks referred to as *modules.* The nodes in each module are densely connected to each other while being sparsely connected to other nodes of the brain, forming a *community structure (missing citation)*. In this context, the brain must balance its capacity to segregate information within modules to perform a specialized neural function in local circuits, with its ability to integrate information between modules. The functional organization of the brain into modules and their functional integration and segregation are data-driven based on the FC matrix between all nodes (cortical regions of Interest [ROIs] in this context). Functional segregation tells us how well the nodes within the modules (cortical ROIs in this context) tend to have a statistical dependence between their time-series [Sporns, 2013]. Contrarily, functional integration represents the joint processing of specialized information across subnetworks (or *modules*). In the context of our study, we could posit that during the postural perturbation, brain sub-networks or modules must perform segregated functional tasks such as processing visuo-sensory inputs, planning for the compensatory response, and actually executing the motor response, while maintaining a certain level of sensorimotor integration between modules to execute a timely, well-measured and coordinated response. To better understand the applications of graph theory in neuroscience, we recommend a review by [Sporns, 2013].

However, to better understand the dynamic aspects of postural control, one needs to study the association between the posturography/balance control measures and the cortical activity recorded during the motor task. The relevant literature on neuroimaging of postural control in a pediatric TBI population is found in [Diez et al., 2017] where the postural control indices were reported to be correlated with increased connectivity in the prefrontal resting-state FC. In another study by [Wang et al., 2016], the postural control change and age were shown to have significant interaction with the prefrontal and sensorimotor connectivity, thus indicating the differences in connectivity across age populations. Although these studies demonstrate the correlation between the EEG sensor-based FC and postural control [Varghese et al., 2019]; [Edwards et al., 2018], sensor-space FC must be interpreted with caution because of its spurious nature associated with the inherent challenge caused by volume conduction; instead, the source-space EEG is recommended for measuring the FC [Bastos and Schoffelen, 2015]; [Handiru et al., 2018]; [de Steen et al., 2016].

For more comprehensive literature on the neuroimaging of human postural control and balance function, readers may refer to the systematic meta-analysis review article [Wittenberg et al., 2017]. As most of the afore-mentioned articles focus only on healthy individuals (young and elderly), the inferences do not offer a complete understanding of balance deficits in the clinical populations suffering from such as brain injury and stroke. Moreover, the fMRI-based neuroimaging of the postural task is impractical due to the device constraints of an MRI scanner; thus EEG offers a unique advantage of being able to noninvasively record the neural recordings while the participant is performing the postural task.

In a preliminary study published by our group, we showed that the N1 amplitude (a type of event-related negative potential) and the center of pressure (COP) displacement were lower in the TBI group as compared to healthy controls [Allexandre et al., 2019]. Further we would like to investigate the changes in cortical connectivity due to postural perturbation in TBI and healthy individuals. We hypothesize that the postural control deficit in TBI could be associated with the altered FC during a balance perturbation task.

While several studies investigated the changes in the cortical activity during the lower-limb motor tasks [Slobounov et al., 2013]; [Munia et al., 2017]; [Wittenberg et al., 2017], the current understanding is still inadequate. Not only should we be able to quantify the functional measures using kinematic factors (e.g., trajectory, the center of pressure, limits of stability, etc.) but also be able to complement the underlying neurophysiological mechanism of balance control through multimodal information. Thus, in this article, we present a group-level analysis of structural and functional mechanisms of postural instability in TBI patients.

Notwithstanding the published literature on functional connectivity changes during postural control tasks in healthy individuals [Varghese et al., 2019]; [Wang et al., 2016], the studies are scarce in TBI or other clinical populations. This motivated us to bridge the knowledge gap of the neural correlates of balance deficits in TBI.

The current understanding points to the fact that TBI can cause both focal injuries and diffuse axonal injury (DAI), which disconnects large-scale brain networks [Sharp et al., 2014]; [Ham and Sharp, 2012]. DAI is commonly induced by a sudden acceleration-deceleration or rotational force, which is mainly affecting WM networks such as corpus callosum and subcortical WM structures [Adams et al., 1989]; [Basser and Pierpaoli, 1996]; [Inglese et al., 2005]. The injuries are seldom visible on computed tomography (CT) and conventional MRI scans, besides the fact that not many approaches are available to assess the WM damage [Adams et al., 1985]. Previous studies have suggested that the DAI is only visible by using advanced neuroimaging techniques that can distinguish microstructural tissue damages [Smith et al., 2019]; [Douglas et al., 2018]. Using Diffusion tensor imaging (DTI), WM connectivity, and integrity changes can be identified by several measures of diffusion anisotropy along fiber tracts [Douglas et al., 2015]. Fractional anisotropy (FA) measures the directionality of water diffusion or the amount of diffusion asymmetry within a voxel or WM tract. Mean diffusivity (MD) quantifies the total water diffusion, regardless of its direction [Uddin et al., 2019]. Mode of anisotropy (MA) is a measure recently developed to help differentiate the local diffusion profile between a planar disc-like (as in fiber crossing) shape and a linear, cylindrical shape (as in unidirectional converging fibers bundle)[Ennis and Kindlmann, 2005].

DTI approach has been quite promising in evaluating WM DAI. Previous studies have shown that damaged WM due to TBI causes increased MD and reduced FA in WM [Shenton et al., 2012]. Our previous work reported the relationship between region-based DTI connectivity (FA values) and physical and cognitive performances in TBI patients [Alivar et al., 2020]. Although a few studies have explored the association between motor impairments and WM integrity in TBI populations [Caeyenberghs et al., 2010]; [Caeyenberghs et al., 2011]; [Drijkoningen et al., 2015], we would like to further investigate the association between the changes in DTI-based measures and balance deficit-related FC in TBI.

This work aims to study the graph-theoretical properties in an FC graph computed from the source-constructed EEG during a balance perturbation task in TBI. This is the first study to report the changes in functional integration and segregation during a postural control task in TBI to the best of our knowledge. Another key contribution is the investigation of the association between the structural and functional connectivity pertaining to postural control in TBI and its relation to balance outcomes.

## Materials and Methods

### I. Participants

In this study, 18 individuals with chronic TBI and 18 healthy controls (HC) with no history of brain injury participated. Participants were recruited through our in-house patient information management system and Kessler Institute for Rehabilitation. Upon careful visual inspection of the raw EEG data, we excluded the data from 4 participants (1 TBI, 3 HC) as the signals were too noisy. This resulted in the inclusion of 17 TBI and 15 HC subjects in our EEG data analysis. Our DTI analysis was further limited to a subset sample of 12 TBI and 9 HC because they failed to meet MRI related inclusion/exclusion criteria or refused to consent to the MRI scan. A summary of the demographics is presented in Table 1. Inclusion criteria for participants are they must be- 1) aged between 18-65 years, (2) diagnosed with a TBI at least 6 months ago (or be a healthy control), (3) medically stable in the past 3 months, (4) be able to stand unsupported for 5 minutes. Exclusion criteria are as follows- (1) history of lower-limb injury in the past 90 days, (2) history of medication that could affect the balance/muscle coordination, (3) have any additional orthopedic, neuromuscular, or neurological conditions that could affect balance (4) have a penetrating TBI, and (5) have a history of previously diagnosed balance impairments (prior to TBI).

**Table 1:** Summary of Participant Characteristics.

| Parameters | TBI | HC | p-value |
| --- | --- | --- | --- |
| N | 17 | 15 |  |
| Severity | Mild = 3; Moderate = 3;<br>Moderate/Severe = 3; Severe = 8 | NA | NA |
| Years post injury (mean, range) | 10.04, [1.67,57.87] | NA | NA |
| Age (mean ± std) | 48.7 ± 12.5 | 47 ± 12.8 | 0.7 |
| Gender (M/F) | 13/4 | 8/7 |  |
| Height in cm (mean ± std) | 177.3 ± 8.6 | 171.9 ± 10.8 | 0.13 |
| Weight in kg (mean ± std) | 90.87 ± 22.8 | 78.26 ± 19.4 | 0.11 |

### 2. Data Acquisition

#### (A) MRI Data Acquisition and Processing

A high-resolution T1-weighted image using MPRAGE sequence was acquired using the Siemens Skyra 3T scanner (Erlangen, Germany) at Rocco Ortenzio Neuroimaging Center at Kessler Foundation. The following settings were used in the whole-brain volumetric acquisition with the following specifications: 1-mm isotropic voxel resolution, TE=3 ms, TR=2300 ms, 1-mm thick 176 slices, Field of View (FOV) 256×256 mm^2^.

A diffusion-weighted (DW) dataset were collected with 64 noncollinear DW gradients with b=1100 s/mm^2^ and eight b=0 images, matrix size=128×128, FOV=256×256 mm^2^, TE=75 ms, TR=9000 ms, isotropic voxel dimensions=2mm, and 66 slices. The gradient-echo field map images were acquired to correct for geometrical distortion caused by susceptibility artifacts. The image processing was performed using the FSL 6.0 toolbox that includes: 1) skull stripping using the Brain Extraction Tool (BET), 2) eddy current correction, 3) DTI-FIT diffusion tensors using FDT (FMRIB’s Diffusion Toolbox, http://www.fmrib.ox.ac.uk/fsl, Oxford, UK). The generated results include parametric maps of DTI metrics, including mean diffusivity(MD), mode of anisotropy (MA), and fractional anisotropy (FA). Finally, the global FA, MD, and MA values for each subject were obtained by averaging the corresponding FA, MD, and MA values of the voxels in the skull-stripped mask, using FSL maths [Smith et al., 2004]. Also, the voxel-wise statistical analysis of the FA images was carried out using the Tract-based spatial statistics (TBSS) [Smith et al., 2006] tool from FSL. First, all the FA data across subjects were aligned to the high-resolution FMRIB58 FA standard space image using the nonlinear registration [Andersson et al., 2007]; [Smith et al., 2007], and subsequently mapped into a 1×1×1mm standard space. These data were then averaged across subjects to obtain the group’s mean FA image, from which the mean FA skeleton was derived as a reflection of the center of fiber bundles. Finally, the subject-specific FA images were projected onto the mean FA skeleton to perform a between-group analysis with the FSL function “randomize” and the threshold-free cluster enhancement (TFCE). In addition to FA, diffusivity maps based on MD and MA were created as described above.

#### (B) Postural Data Acquisition and Processing

A computerized dynamic posturography (CDP) platform (NeuroCom Balance Master, NeuroCom Intl, Clackamas OR) as shown in Fig. 2 was used to study the neural and postural response to unpredictable balance perturbations. Henceforth, we use the term “CDP platform” and “balance platform” interchangeably. The participants were asked to stand on the balance platform and they were subject to five blocks of random, unpredictable perturbations in the anterior (forward) and posterior (backward) sinusoidal perturbations at 0.5Hz, (two cycles for a total of 4s duration) at low (0.5cm) and high (2cm) amplitude with an inter-trial separation of 4-8s. Thus, there are 2×2 combinations of perturbation anterior and posterior (direction), low and high (amplitude). We measured the center of pressure (COP) time series from the ground reaction force data. The COP data were epoched, low-pass filtered (10Hz), detrended, and averaged across trials and conditions for each subject. The COP displacement was computed as the cumulative distance traveled by the trial average COP in the anterior/posterior direction for the first 2s of the balance perturbation. In this article, our analysis is limited to the high amplitude perturbation in the posterior direction, which led to greater instability across all subjects.

**Figure 2:**
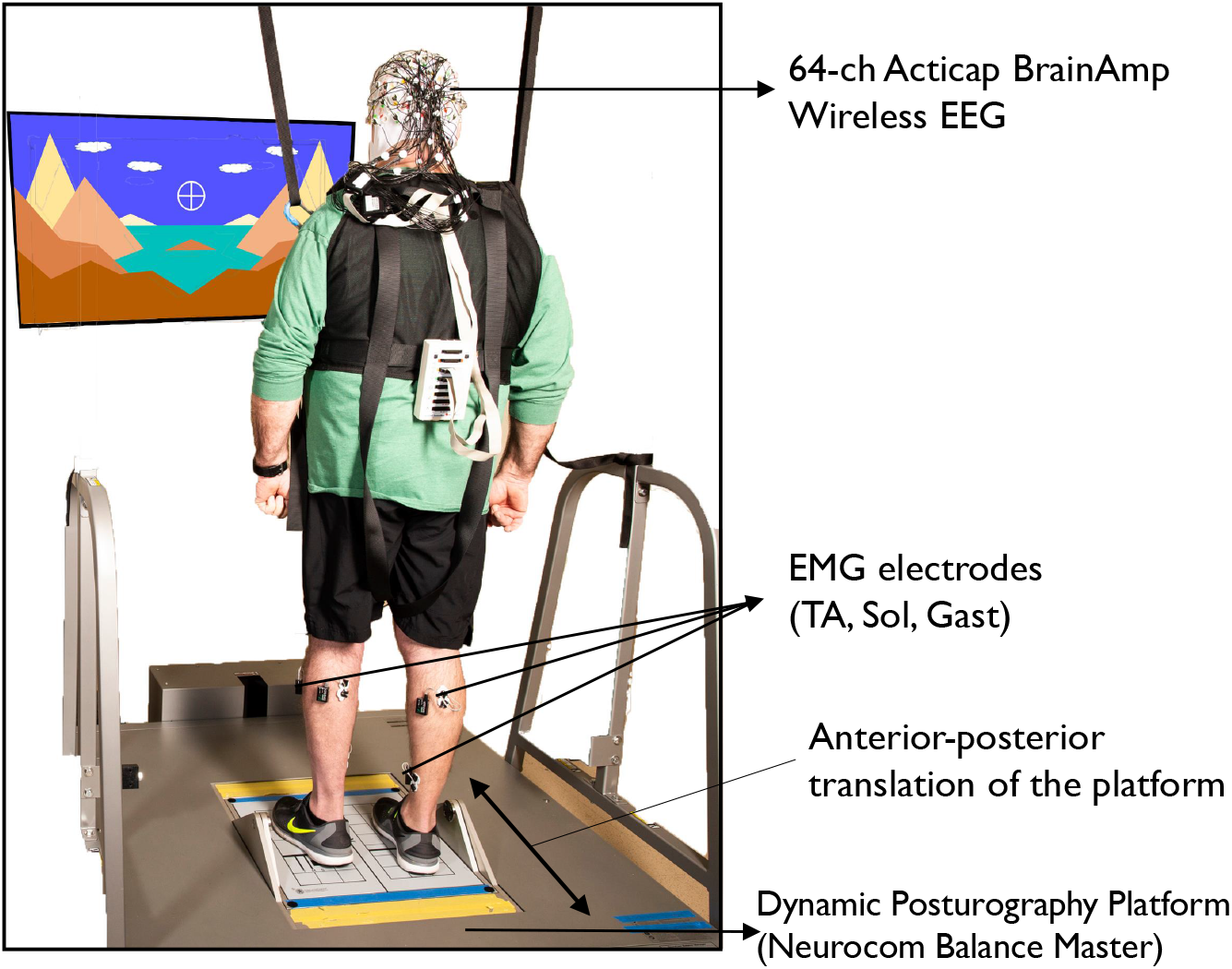
A representative subject standing on the dynamic posturography platform used in the study

#### (C) EEG Data Acquisition

Throughout the duration of the task, the brain activity was recorded using the 64-channel EEG system (ActiCAP BrainAmp standard, Brain Products®, Munich, Germany) positioned according to the International 10-20 systems where the electrode änd “AFz” serve as the common reference and ground respectively@”FCzänd “AFz” serve as the common reference and ground respectively. EEG data were sampled at 500Hz and the skin-electrode contact impedance was ensured to be below 20k ohms by applying electrode gel. The EEG electrode positions were 3D digitized using the Brainsight® navigation system.

### 3. Data processing

#### (A) EEG Data preprocessing

EEG recordings were analyzed offline using the EEGLAB toolbox [Delorme and Makeig, 2004]. The raw EEG signals were first downsampled to 250Hz, followed by band-pass filtering between 1Hz and 50Hz using a 4^th^ order Butterworth filter. Thereafter, the line noise was removed using the Cleanline plugin for EEGLAB; artifact-contaminated channels and noisy continuous-time segments were removed using the Artifact Subspace Reconstruction (ASR) plugin for EEGLAB [Mullen et al., 2015]. The ASR parameter to reconstruct the high variance subspace was set as 20 based on the findings reported in [Chang et al., 2020]. Subsequently, we performed the Independent Component Analysis (ICA) via extended Infomax algorithm [Makeig et al., 1996]. The resulting ICs presumably originated from the following physiological or miscellaneous sources—brain, muscle, eye, line noise, channel noise, heart, or other—were labeled accordingly with the ICLabel plugin in the EEGLAB toolbox [Pion-Tonachini et al., 2019]. The ICLabel is a machine learning approach, which has been trained to classify the ICs derived from EEG data based on several characteristics such as spectral properties, brain topography. Furthermore, to validate the ICs supposedly generated by neural sources, the DIPFIT tool (available in EEGLAB) was used that localizes the dipoles within the brain volume [Delorme et al., 2012]; [Oostenveld and Oostendorp, 2002]. In our study, we only retained the ICs that were (1) identified as ‘Brain’ by the ICLabel plugin and (2) having a residual variance of <15% after localization by DIPFIT modeling. Although we used the ICLabel for automating the labeling of ICs, we also visually inspected whether the characteristic topography and spectral properties of ICs corroborated with their respective labels. Once the ICs were selected, we used the back-projected sensor EEG for source localization. The EEG data processing pipeline is summarized in the block diagram shown in Fig. 3.

**Figure 3:**
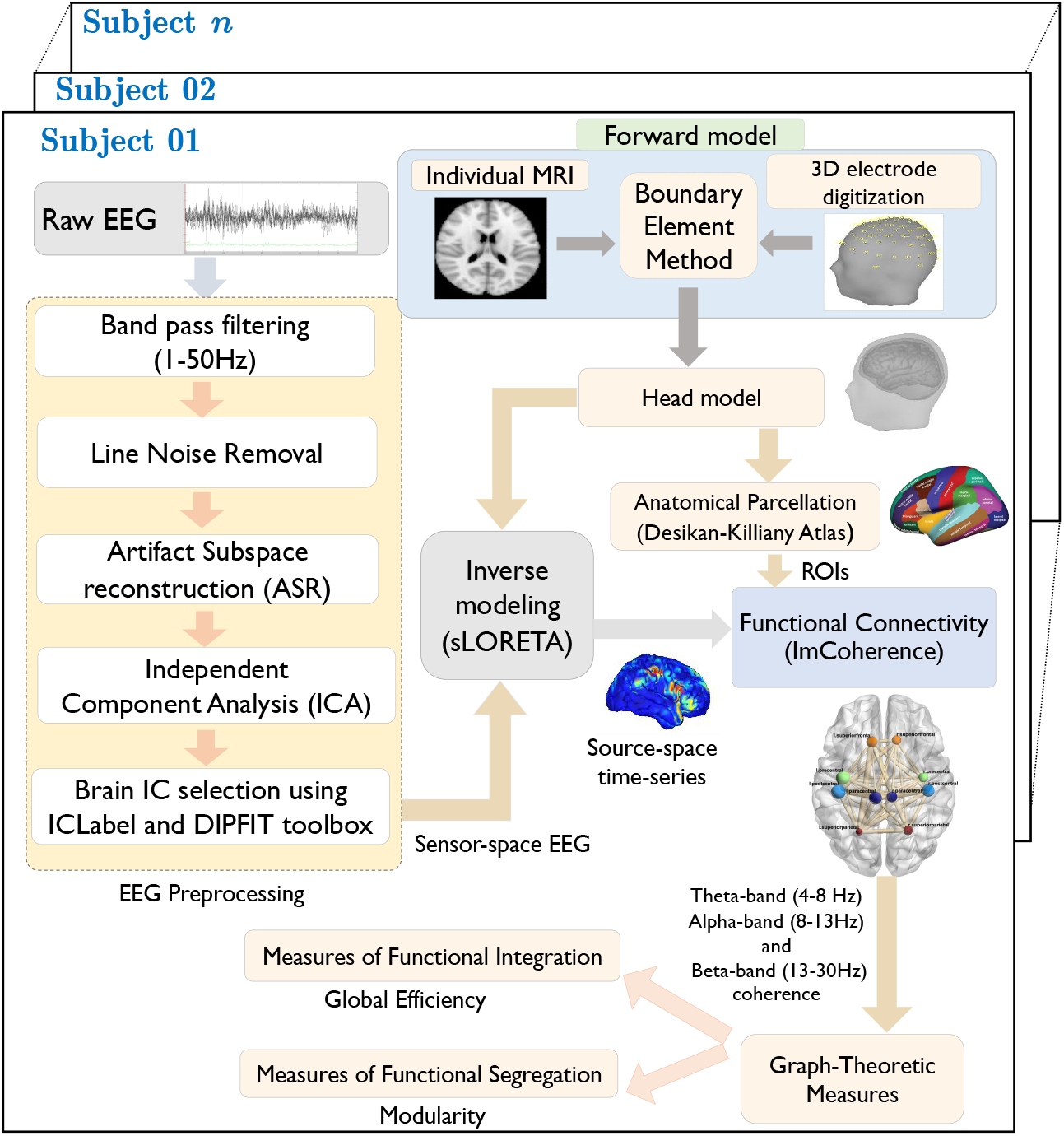
Block diagram of the EEG processing pipeline

#### (B) EEG Source localization and Functional Connectivity Estimation

EEG Source Localization (ESL) allows one to study the cortical dynamics using the underlying cortical sources estimated from the sensor-space EEG [Michel and Brunet, 2019]; [Handiru et al., 2018]. As shown in Fig. 3, ESL is implemented in two stages: forward and inverse modeling. Forward modeling is done to compute the volume conduction model (VCE), which realistically approximates the electromagnetic field propagation through different layers in the head model (scalp, skull, cerebrospinal fluid, grey matter, and WM). Using the subject-specific anatomical data comprising of T1-weighted MRI ad 3D digitized EEG positions, a realistic forward model is created using the Boundary Element Method (BEM) [Fuchs et al., 1998]; [Gramfort et al., 2010]. Inverse modeling based on sLORETA (standardized Low-Resolution Electromagnetic Tomography) algorithm [Pascual-Marqui, 2002] is used to compute the cortical time series data. More mathematical details of the ESL can be found in the supplementary material (Section S1). All the aforementioned steps were implemented using Brainstorm software [Tadel et al., 2011].

Once the source time series is obtained, we computed the region-of-interest (ROI) time series by averaging the time-series of all the voxels with each anatomical region parcellated according to the Desikan-Killiany Atlas. Between each ROI, we then estimated the functional connectivity using imaginary coherence 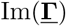 with 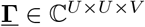 being a three-way tensor, where *U* and *V* are the number of ROIs and frequency bins of interest, respectively. In this study, 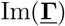 is computed during the *baseline* (*t* = −2s to 0s) and *task* (*t* = 0s to 2s) period for each frequency bin and averaged across frequencies to obtain coherence measures for the theta band (4-8Hz), alpha-band (8-13Hz) and beta-band (13-30Hz), given their distinct role in sensorimotor function [Omlor et al., 2011]; [Buchholz et al., 2014].

#### (C) Graph-Theoretic measures

As mentioned earlier, the *functional segregation* informs about the local network properties, such as how densely connected are the sub-networks (or *modules*) [Newman, 2006] (Fig. 1A). On the other hand, *functional integration* captures the information about the ease of interaction between these sub-networks (Fig. 1B). The following graph-theoretic measures to characterize the functional segregation and integration of the network were computed for theta-, alpha- and beta-band connectivity. The weighted rather than the binary definition of the connection or edge between nodes are used, which is defined as the normalized connectivity strength with values between 0 and 1 [Rubinov and Sporns, 2010].

##### The Measure of Functional Integration

###### Global Efficiency (*GE*)

It describes how well connected is the neighborhood of a node. It is a measure of the efficiency of information transfer among all pairs of nodes in a graph, given by

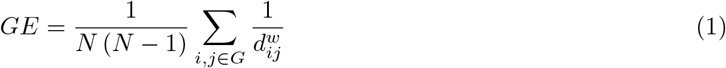

where, *N* denotes the number of nodes in a graph *G* and 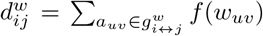 denotes the shortest weighted path length between the nodes *i* and *j* in a graph *G* computed using the Djikstra’s algorithm. 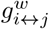 corresponds to the shortest weighted path between the nodes *i* and and *j*, and *f* (*w_uv_*) denotes the mapping from connection weight to the path length (i.e., *f* (*w_uv_*) = (*w_uv_*)^−1^). In the context of brain connectivity, *w_uv_* is the imaginary coherence value measured between the nodes *u* and and *v*. Greater GE would reflect overall more direct communication i.e., shorter path length between nodes.

##### The Measure of Functional Segregation

*Modularity (M)* is a measure of functional segregation that describes the *community structure* of a network [Newman, 2006]. It compares the number of connections within *modules (or sub-networks*) to the number of connections across *modules.* Although there are different ways to compute the modularity, here we used Newman’s algorithm as provided in the BCT toolbox [Rubinov and Sporns, 2010]. The modularity score *Q^w^* for a weighted connection matrix can be computed as below:

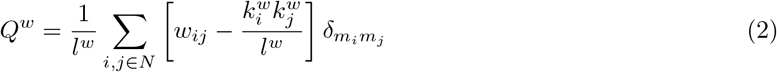

where, *l^w^* is the sum of all weights in the given network (i.e., *l^w^ = Σ_i,j_ w_ij_*), also defined as the Network Strength (*NS*). 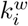 and 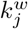 denote the weighted node degree of node *i* and *j,* respectively (i.e., 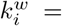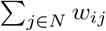). *m_i_* and *m_j_* define the module containing the node *i* and *j* respectively. δ(*m_i_*, *m_j_*) defines the community structure i.e., 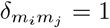, if both nodes i and j belong to the same community.

### 4. Statistical Analysis

The normality assumption for the data distribution was verified using visual inspection and Shapiro-Wilk test with a significance level of *α* = 0.05. We performed the two-sample *t-*test to compare the demographics characteristics (age, height, and weight) and the COP displacement. As the normality assumption was not met, we performed a non-parametric statistical test (Wilcoxon ranksum test) for comparing the Berg Balance Scale (BBS) across groups. To investigate the cortical regions involved during the balance perturbation task, we performed a parametric statistical test (student’s *t*-test) at the population level (TBI and HC) to identify significant voxel differences between the task period (*t* = 0s to 2s) and baseline period (*t* = −2s to 0s). Because of the multiple comparisons, the correction for False Discovery Rate (FDR) was done using the Benjamini-Hochberg procedure with the conservative significance value set at *α*=0.01.

Each imaginary coherence value between any two nodes (or ROIs) was tested for its statistical significance using Schelter’s approach [Schelter et al., 2006] as implemented in the Brainstorm toolbox.

Since the data distribution turned out to be either normal or pseudo-normal, we performed a two-way repeated-measures analysis of variance (2×2 rmANOVA) using a general linear model, to compare the graph metrics across groups (TBI vs. HC) and time (*baseline* period between −2s and 0s and balance perturbation *task* period between 0 to 2s) for each frequency band. Whenever an overall significant effect was found, corresponding post-hoc analyses using a two-tailed *t*-test were performed to examine between-group differences at baseline and/or during the task, between-group changes over time, and/or changes within the group.

The effect sizes are reported using cohen’s D (for *t*-test) and partial eta-squared (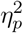 for rmANOVA) values. The statistical analyses related to ANOVA and *t*-test were performed with SPSS (version 26, IBM, NY, USA), with the significance level set at α < 0.05.

Furthermore, to determine whether network measures are associated with the behavioral measures, we calculated the Pearson correlation. In the case of correlation analysis, we identified the influential data points using a commonly accepted criterion of cook’s distance > 4/N [Cook, 1977], where, N is the number of data points in the regression model. The corrections for multiple correlations were done using the Bonferroni correction.

## Results

### 1. Participants characteristics and balance outcomes

A two-tailed *t*-test revealed no significant differences in the baseline demographics (Age, Height, and Weight) characteristics, as shown in Table 1. With regard to the functional outcome measures, as shown in Fig. 4, TBI group showed a significantly (*t* = 3.07, *p* = 0.004, cohen’s D = 1.09) larger COP displacement (mean ± SD = 11.64 ± 4.28, 95% CI =[4.79, 19.18]) as compared to healthy controls (HC) (mean ± SD = 7.82 ± 2.33, 95% CI = [4.7, 13.13]). Due to the non-normal distribution of the BBS, we ran a Wilcoxon ranksum test to compare the BBS of both groups. We observed a significantly lower BBS (*p* = 0.007, z = 2.68) in TBI (median, range = 51, [34,56]) than HC (median = 56, range = [55,56]).

**Figure 4:**
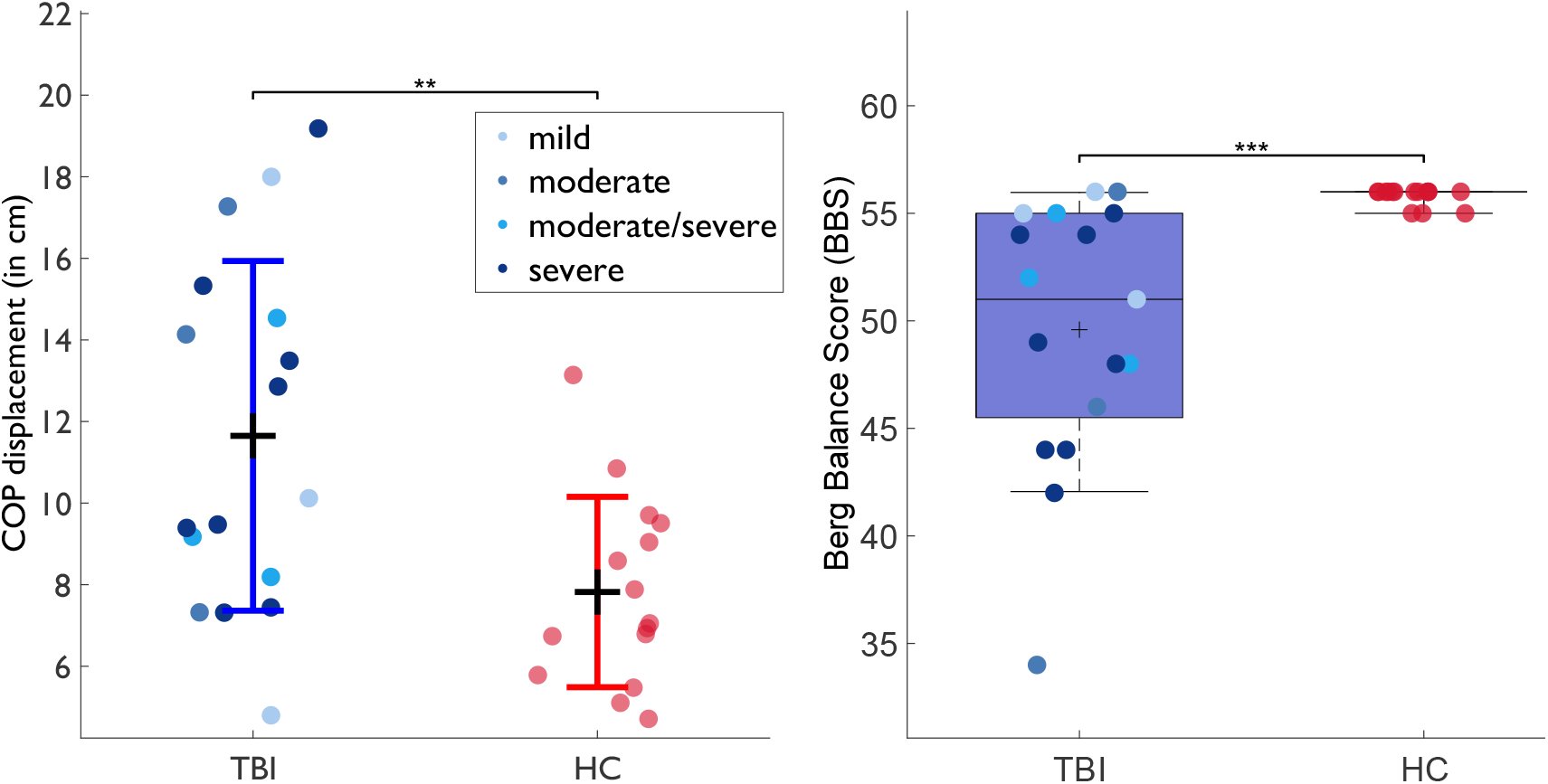
A group-level comparison of COP displacement (in cm) shown on the left. The black horizontal line on the COP plot marks the mean and the colored horizontal line marks the standard deviation. A group-level comparison of the Berg Balance Scale (BBS) is shown on the right as a boxplot due to its non-normal distribution. The horizontal line marks the median. The lower- and upper-hinge of the boxplot corresponds to the 25th and 75th quartile respectively. Statistical significance values are plotted as ***(*p*<0.005), **(*p*<0.01), *(*p*<0.05) respectively.

### 2. Group differences in task-induced cortical activity during balance perturbation

Table 2 lists the key regions of interest that showed a significant activity (EEG source power) during the task as compared to the baseline with the respective MNI coordinates as well as the number of significant voxels within each group. The spatial distributions of the significant voxels in different frequency bands are shown in Fig. 5.

**Table 2:** MNI coordinates for contrast: perturbation task vs. baseline period.

| Region Labels | TBI group |  |  |  |  | HC group |  |  |  |  |
| --- | --- | --- | --- | --- | --- | --- | --- | --- | --- | --- |
|  | MNI coordinates |  |  |  |  | MNI coordinates |  |  |  |  |
|  | x | y | z | t-value | #voxels | x | y | z | t-value | #voxels |
| cingulate gyrus (Left) | -5 | 3 | 93 | 6.06 | 36 | 15 | -1 | 75 | 12.0 | 114 |
| cingulate gyrus (Right) | 0 | -6 | 94 | 5.70 | 36 | 2 | -9 | 92 | 13.45 | 128 |
| paracentral lobule (Left) | -11 | 0 | 119 | 5.92 | 27 | 19 | 8 | 124 | 10.26 | 47 |
| paracentral lobule (Right) | 10 | -4 | 92 | 4.86 | 25 | 12 | -2 | 89 | 8.87 | 61 |
| post-central gyrus (Right) | -14 | -14 | 119 | 4.94 | 29 | 7 | -6 | 122 | 8.91 | 92 |
| pre-central gyrus (Left) | -6 | 4 | 122 | 5.55 | 21 | 12 | 12 | 124 | 8.42 | 68 |
| superior frontal gyrus (Left) | 30 | 1 | 117 | 4.57 | 17 | 23 | 8 | 120 | 12.04 | 125 |
| superior parietal gyrus (Left) | -23 | 22 | 110 | 5.12 | 25 | -30 | 14 | 118 | 5.57 | 52 |
| superior parietal gyrus (Right) | -27 | -17 | 112 | 6.21 | 34 | -20 | -26 | 122 | 9.5 | 67 |

**Figure 5:**
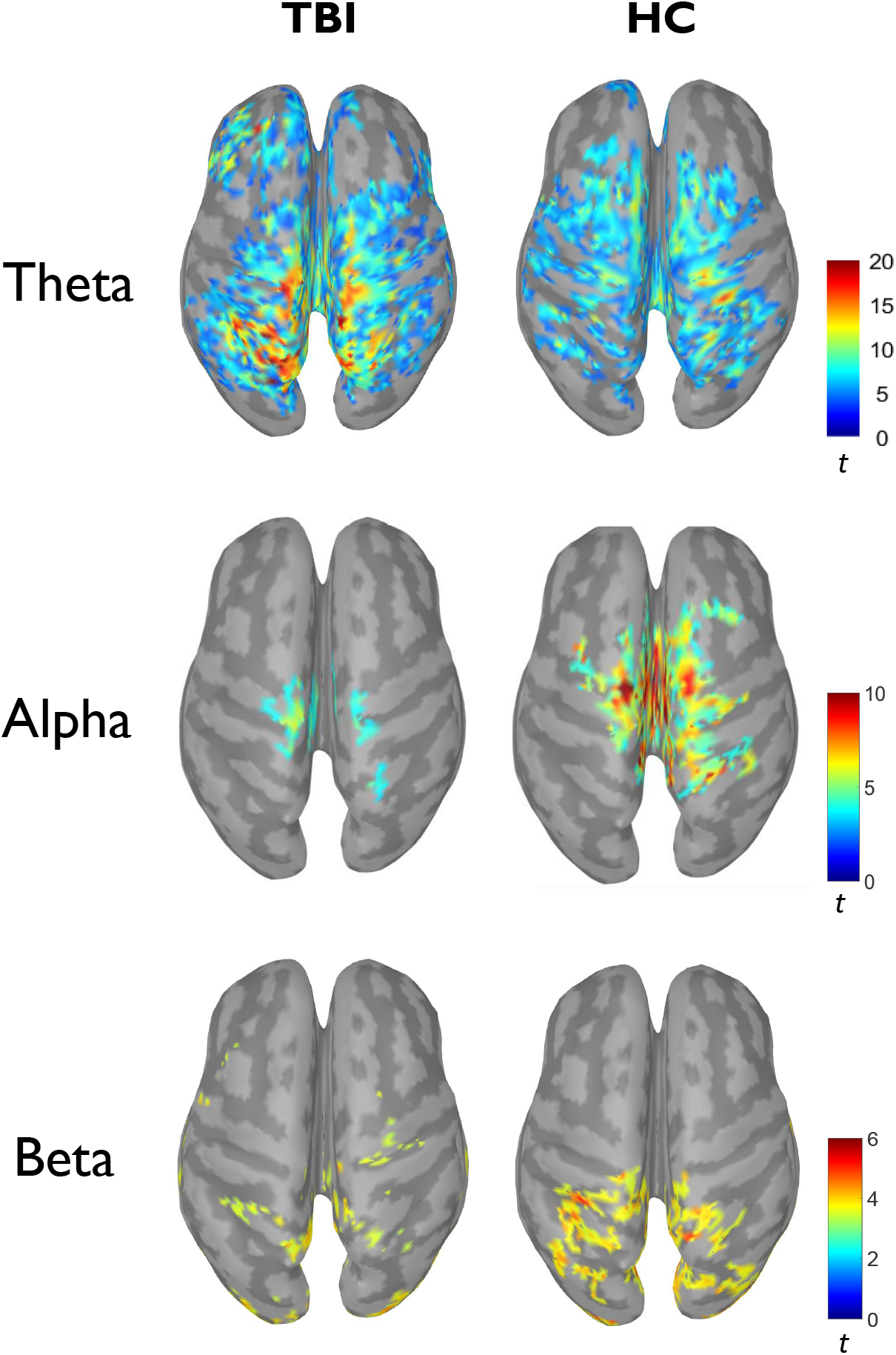
Spatial distribution of task-specific significant voxels mapped onto the cortex within each group. Parametric student’s t-test is used to identify the voxels that are significantly different from the task period (*t* =0 to 2s) as compared to the baseline period (*t* =−2s to 0s). FDR correction is with the significance level of *α*=0.01.

### 3. Two-way Repeated Measure Analysis of Variance

#### (A) Whole-brain functional connectivity network strength

Fig. 6 shows the *non-normalized* functional connectivity and *non-normalized* node strength for each ROIs (defined as the sum of connectivity values to all other ROIs i.e., how strongly connected that ROI is) based on source-space EEG coherence in the theta- and alpha-bands during the baseline period as well as during the balance perturbation task in TBI and HC.

**Figure 6:**
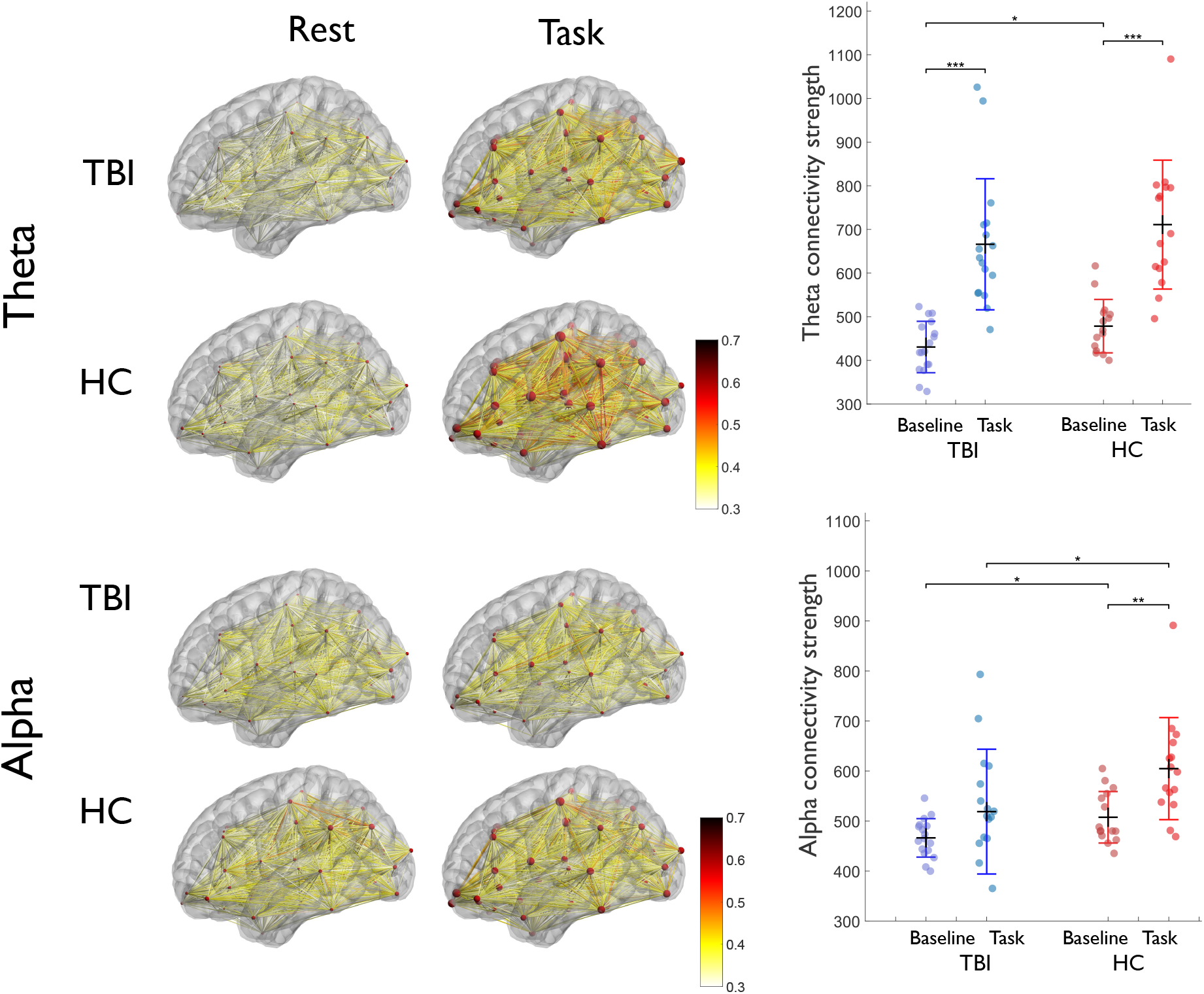
Whole-brain functional connectivity based on source-space EEG coherence in different frequency bands in each group (*TBI* and *HC*) across time periods (*baseline* and *task*). Group-level connections are averaged and plotted as an edge between different ROIs anatomically parcellated using Desikan-Killiany Atlas. For better visualization, the connections are thresholded at edge weight = 0.3. Seed voxels of each ROI are indicated as spheres with a radius proportional to their node strength. On the right side, the boxplot comparison of network strengths during the baseline vs. task period is shown for both the groups. Statistically significant differences are highlighted with asterisk * (p < 0.05), ** (p<0.01), *** (p<0.005), and marginally significant difference is highlighted with a † (p<0.1). Visualization of the cortical map is done with the help of BrainNet Viewer [Xia et al., 2013].

The two-way repeated-measure ANOVA (rmANOVA) conducted on the graph measures are summarized in Table 3 and the descriptive statistics in Table 4. Theta coherence-based network strength (NS) showed a main effect of *Time*, but only a marginal non-significant *Group* effect and no *Group × Time* interaction. Within-group two-tailed *t*-test revealed a significant baseline to task change for both the groups (TBI and HC). Between-group contrast analysis revealed a greater network strength in HC compared to TBI at baseline (Fig. 6).

**Table 3:**
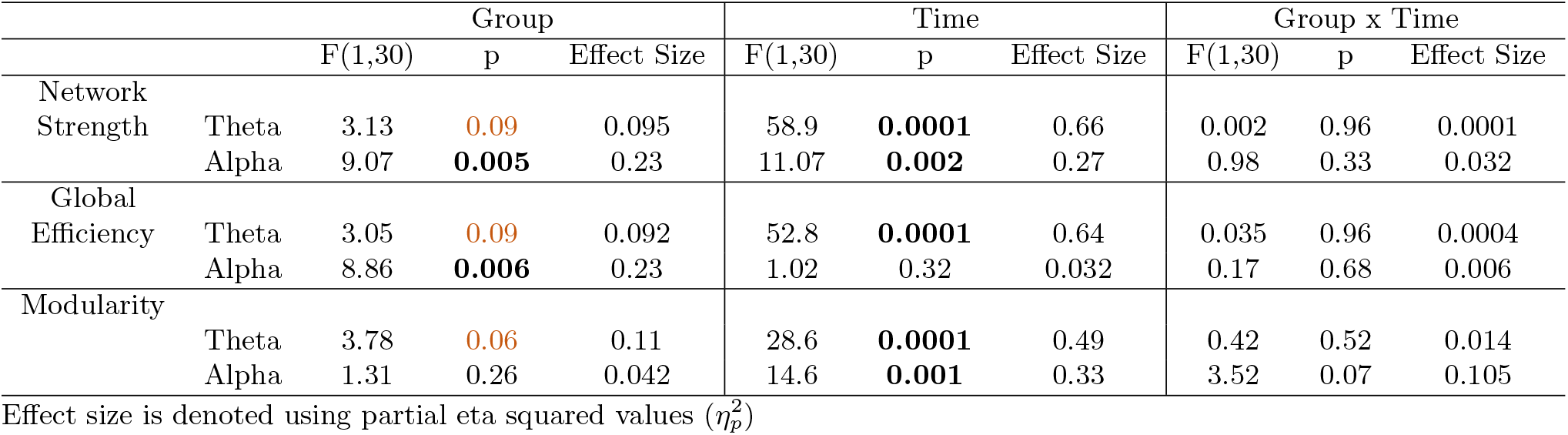
Summary of 2-way repeated measures ANOVA.

**Table 4:**
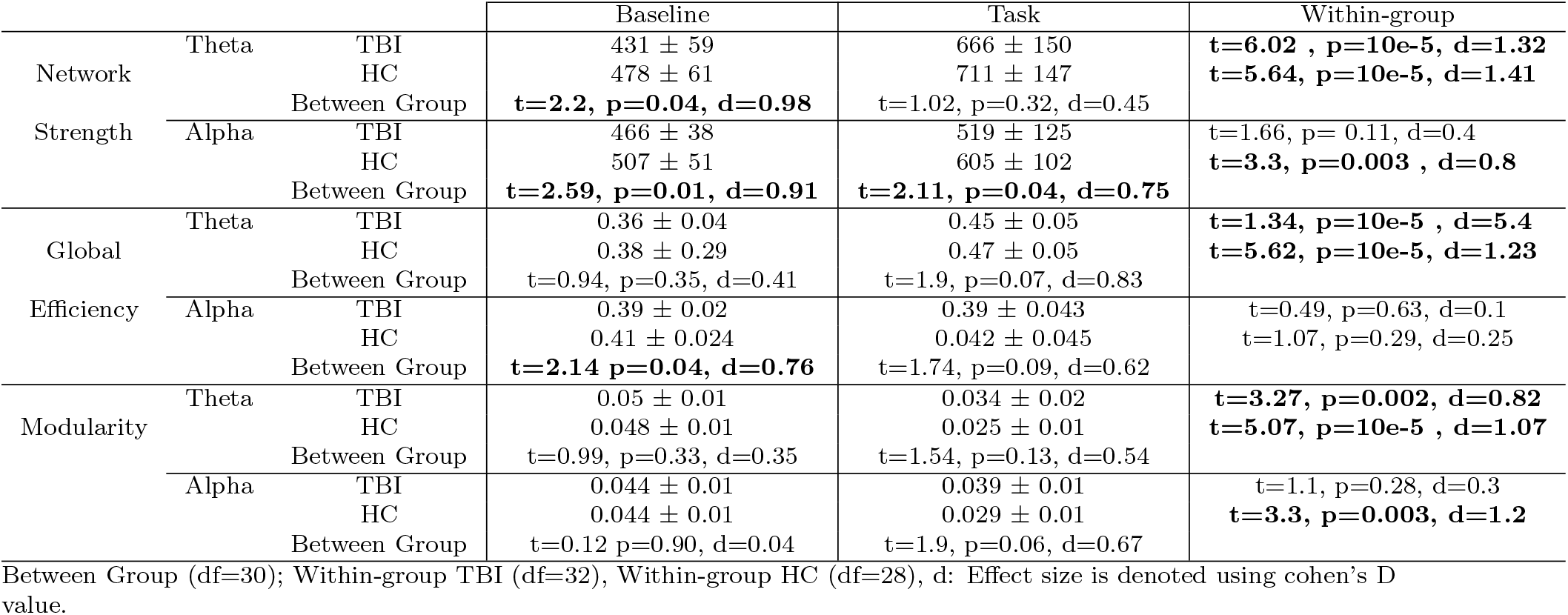
Descriptive statistics and contrast analysis of graph measures.

When comparing the alpha-band network strength, two-way rmANOVA revealed a significant main effect of *Time* as well as *Group*, but no significant *Group × Time* interaction. Between-group contrast analysis revealed a lower network strength (alpha-band) in TBI compared to HC during the task as well as the baseline (Fig. 6).

We did not find any significant differences with the beta-band results, but they are included in supplementary materials (Table. S1) for the sake of completion. Overall, we observed a pattern of globally weaker connectivity and weaker node strength in TBI compared to control.

#### (B) Graph Measures of Functional Integration and Segregation

Global network measures of functional integration and segregation were measured using the Global Efficiency (GE) and Modularity (M), respectively, in different frequency bands (during the *baseline* and balance perturbation *task*) for each group. The boxplot summary of the group-level comparison is shown in Fig. 7. The results for beta-band coherence connectivity were not significant, but for the sake of completeness, they are included in the supplementary material (Table S1 and Table S2).

**Figure 7:**
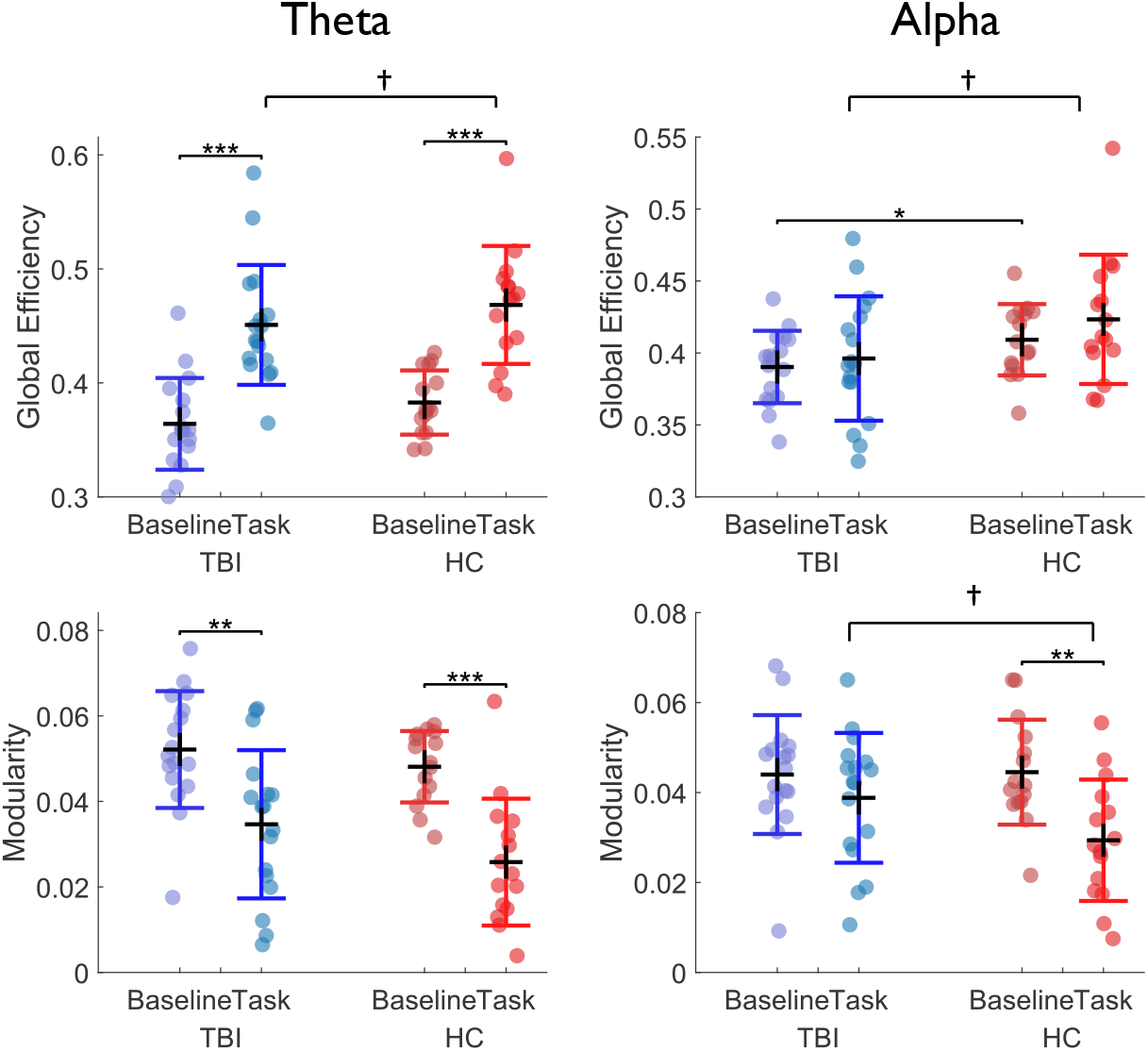
Group-level comparison of network measures of functional integration (top half) and functional segregation (bottom half) across frequency bands. The categorical scatter plot of the network measures during the baseline and during perturbation (task) within each group is shown above. Black horizontal line indicates the mean and the colored horizontal lines indicate the standard deviation. Statistically significant differences are highlighted with asterisk * (p < 0.05), ** (p<0.01), *** (p<0.005), and marginally significant difference is highlighted with a † (p<0.1).

##### Functional Integration

The two-way rmANOVA was conducted on GE independently for each of the frequency bands. Theta coherence-based GE showed a significant main effect of *Time* and a marginal effect of *Group*, but no significant *Group × Time* interaction. As illustrated in Fig. 7, the within-group post-hoc analysis revealed a significant *baseline* to *task* change in GE for both the groups.

In the case of alpha-band GE, the two-way rmANOVA showed a significant main effect of Group but with no significant *Time* effect or *Group × Time* interaction. Moreover, the within-group post-hoc analysis revealed no significant *baseline* to *task* change in GE for either group.

##### Functional Segregation

To investigate the task and group effect on functional segregation, the two-way rmANOVA comparison was performed on the modularity (M) values derived from different frequency bands. We noticed a significant effect of *Time* and *Group* for Theta-band M, but no *Group × Time* interaction. The post-hoc within-group analysis showed a significant *baseline* to *task* decrease within both groups.

In the case of alpha coherence-based M, there was a significant effect of *Time* and a marginally significant *Group × Time* interaction but not of *Group.* Indeed, as shown in Fig. 7, regions are functionally less segregated during the balance perturbation *task* compared to *baseline* for HC but not for TBI. The post-hoc analysis of baseline to task change in alpha modularity revealed a marginally significant difference (p = 0.067) between HC (−0.015 ± 0.013) and TBI (−0.005 ± 0.017). Intuitively this suggests that during the perturbation task, more cortical areas (aka ‘network node’) are interacting with each other in HC, thus reducing the functional segregation, but not in TBI. Given the lack of significance of beta-band results, we have included the results in the supplementary material.

### 4. Group differences in global DTI measures

Changes in structural networks were assessed by performing group comparison of the global measures of DTI as fractional anisotropy (FA), mean diffusivity (MD), and mode of anisotropy (MA) computed for the whole brain for the subset of 12 TBI and 9 HC with DTI data. All three metrics showed significant differences in the structural integrity between TBI and HC subjects (see Fig. 8). More specifically, the global FA measure is significantly lower (F(1,19) = 10.7, *p* = 0.004) in TBI (mean ± SD =0.234 ± 0.015, 95% CI: [0.225,0.243]) than in HC (mean ± SD = 0.256 ± 0.013, 95% CI: [0.247,0.265]). MD values are significantly higher (F (1,19) = 6.68, *p* = 0.018) in TBI (mean ± SD =0.0011 ± 0.00008, 95% CI:[0.0011,0.0012]) than HC group (mean ± SD =0.001 ± 0.00008, 95% CI: [0.001,0.0011]). Also, as shown in Fig. 8, significant difference (F(1,19) = 5.66, *p* = 0.028) was observed for the global MA between TBI (mean ± SD =0.214 ± 0.022, 95% CI: [0.202,0.227]) and HC (mean ± SD =0.236 ± 0.017, 95% CI: [0.225,0.247]).

**Figure 8:**
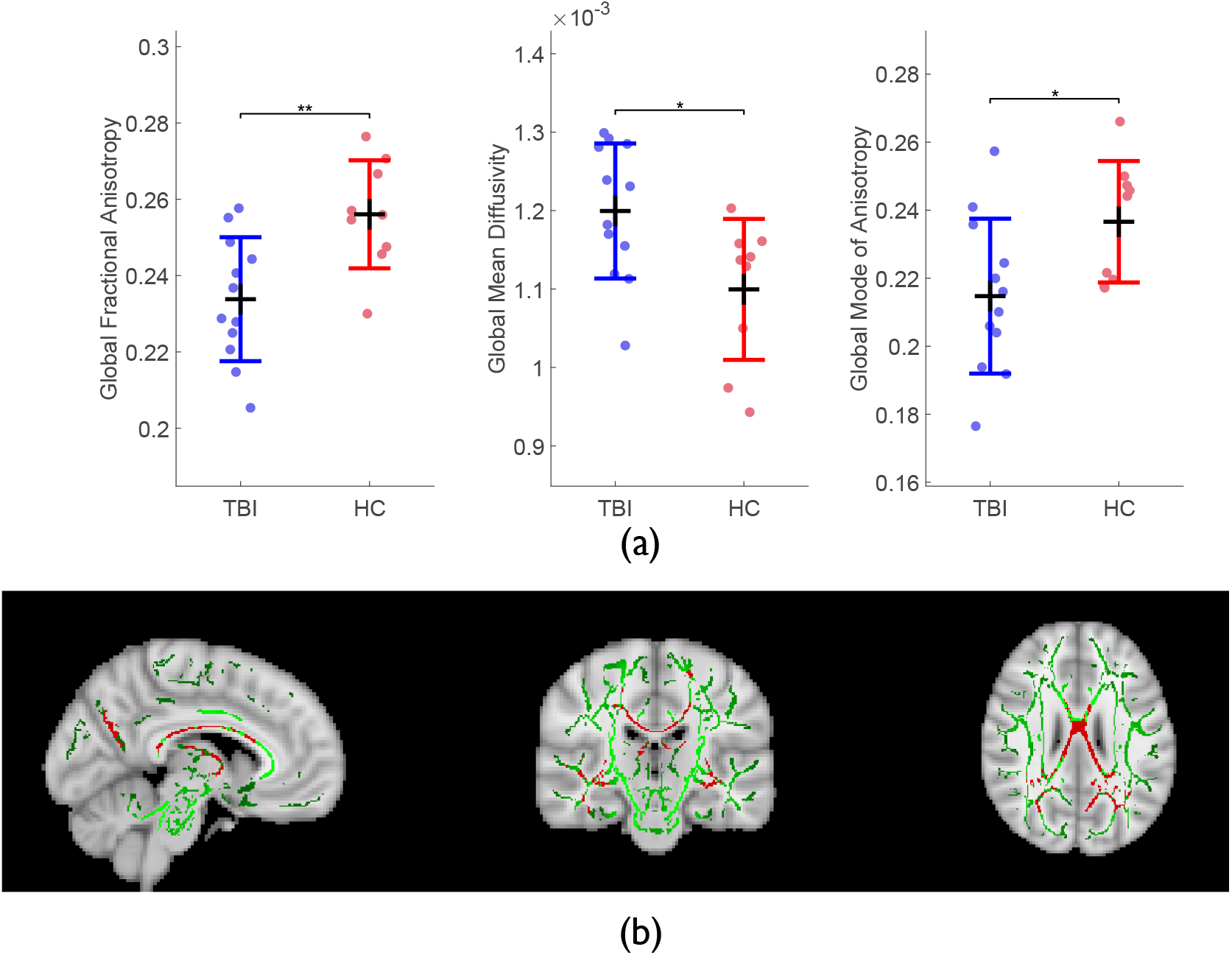
(a) Boxplots showing the differences in global DTI measures across groups. Statistically significant differences are highlighted with *(p<0.05), **(p<0.01). (b) Three views (sagittal, coronal, and axial) of significant differences in FA skeleton between TBI and HC group. The underlying image is the MNI152-T1 template, the green contour is the mean FA skeleton, and the red one shows the regions with significantly higher (p = 0.02) FA values in HC group compared to TBI.

To address the structural changes in brain WM between groups, we conducted the voxel-wise analysis for the FA images using TBSS as shown in Fig. 8(b). Areas in red are regions where FA was significantly higher (*p* = 0.02) in HC than TBI, specifically in Corpus Callosum and Thalamus. No significant differences were observed between groups’ comparison of MD and MA skeletons.

### 5. Association between DTI measures and behavioral measures

To explore the underlying association between WM damage and balance, the correlation between DTI global metrics (FA, MD, and MA) and behavioral measures i.e., COP and BBS, in TBI patients were studied. Given the multiple correlation analyses (3 DTI measures × 2 behavioral measures), we ran the Bonferroni correction with the corrected *p =* 0.05/6 (i.e., 0.008). The only correlation that survived the Bonferroni corrections was a significant negative relationship between MD and BBS (*r* = −0.78, *p* = 0.0043 without and *r* = −0.64, *p* = 0.02 with outlier; Fig. 8 (b)) which is basically implying that higher MD values or more damaged brain tissues (more free diffusions) are correlated with lower BBS.

### 6. Association between EEG graph-theoretic measures and behavioral measures

To explore the brain-behavioral relation, Pearson correlation coefficients between EEG graph measures in each of the frequency bands and behavioral measures (i.e., COP displacement (in cm) and BBS) within the TBI group were calculated. To avoid spurious results given the small sample size, any outlier with a strong influence on the correlation based on the cook’s distance D > 4/N criterion [Cook, 1977] was removed. Both results with and without the outliers are reported for completeness (Fig. 9 (a)). As the correlation analyses involved 3 graph measures (NS, GE, and M) across 3 frequency bands, and 2 behavioral measures (COP and BBS), we did the Bonferroni correction for multiple comparisons with the corrected *p-value* being p= 0.05/18 (i.e., 0.0027). Upon correcting for multiple comparisons, we only observed a significant negative correlation between the Theta Modularity and BBS (*r* = −0.72, p = 0.001).

**Figure 9:**
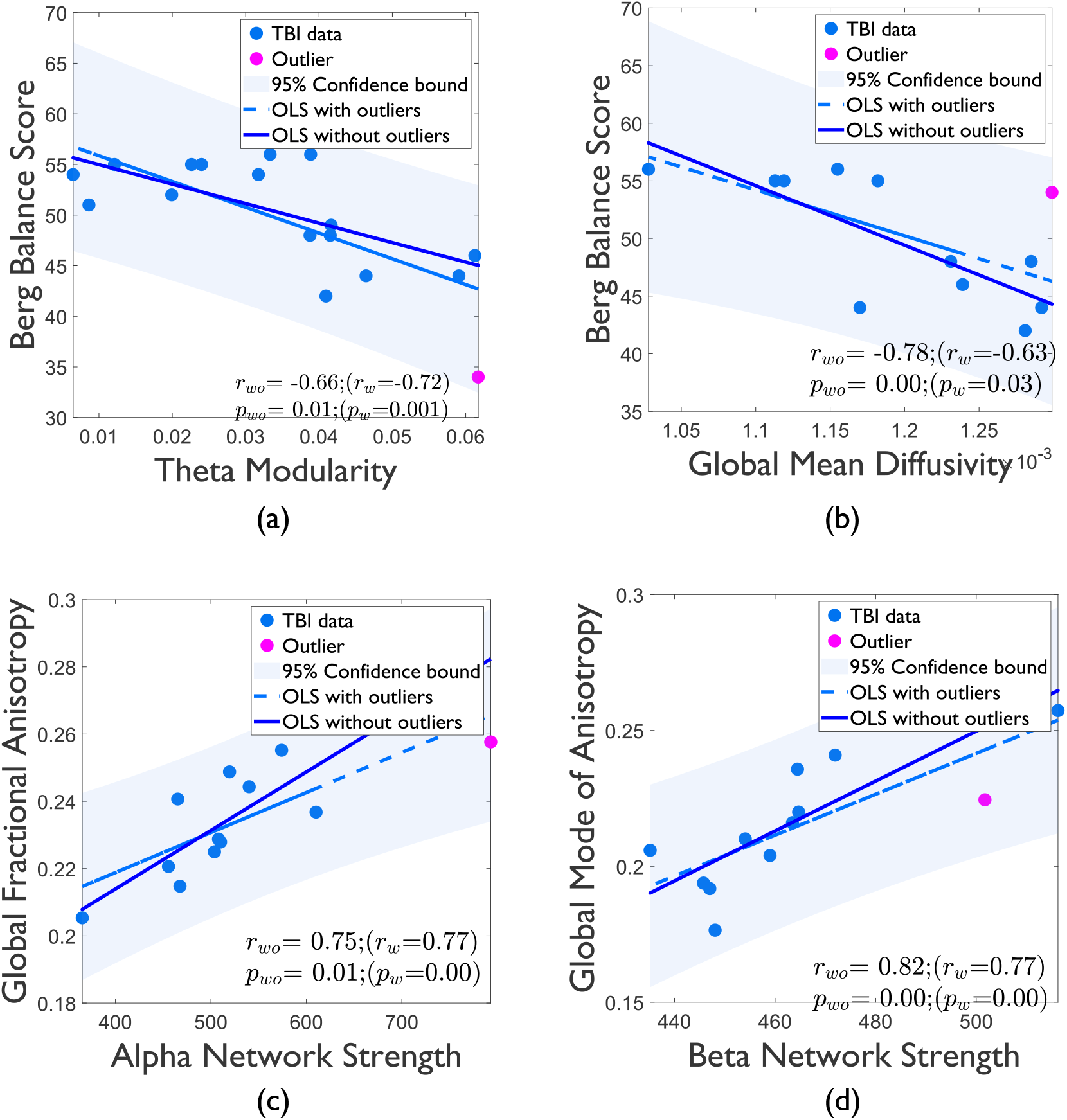
Plots of the ordinary least squares regression correlation between the functional outcome measure (BBS in the Y-axis) and (a) Theta Modularity and (b) Global Mean Diffusivity respectively in the x-axis. (c) The significant correlation between the global fractional anisotropy and alpha-band network strength during the postural control task within the TBI group, and (d) global mode of anisotropy and beta-band network strength. The effect of outliers on the correlation is highlighted with two separate regression lines (*dashed line* when outliers are removed, *solid line* when the outliers are retained). *r* and *p*-values indicate the Pearson correlation coefficients and the significance level, respectively. *r_w_* and *p_w_* denote the correlation statistics when outliers are also taken into account; *r_wo_* and *p_wo_* denote the correlation statistics without outliers

### 7. Association between global DTI measures and global EEG connectivity graph measures

To explore the underlying association between structural and functional connectivity graph metrics during postural control task, the correlation between each of the three functional graph metrics (GE, M, and NS) and global DTI metrics (FA, MD, and MA) were analyzed using the Pearson correlation test. Due to the multiple comparisons (3 DTI measures, 3 graph measures in each of the 3 frequency bands), the Bonferroni-corrected *p*-value is 0.05/27 (0.002). Only the beta-band network strength was positively correlated with the global MA (when outliers were removed: *r* = 0.82, *p* = 0.002; but *r* =0.77, *p* = 0.003 if included). Moreover, the alpha-band network strength was marginally non-significantly correlated with the fractional anisotropy (*r* = 0.77, *p* = 0.003 without and *r* = 0.75, *p* = 0.01 with outliers). Overall these results suggest a relationship between the structural integrity of the white matter system and the strength of the functional connections. For the sake of completion, we report other significant correlations that did not survive the Bonferroni correction in the supplementary material.

## Discussion

In this study, we demonstrated the neural correlates of balance deficits using global and topological measures of functional connectivity and structural integrity of the WM in the TBI population as compared to healthy controls. To the best of our knowledge, this is the first report of EEG-based functional connectivity measures during a postural perturbation task in TBI and its association with the altered WM integrity.

### 1. Effect of TBI on balance

The widely used clinical measure of balance function using the BBS showed an overall balance deficit and a significantly lower BBS score in the TBI population compared to HC. However, there was a wide range of deficits with half of our sample scoring near the healthy maximal values of 56, while the other half below 50, with the knowledge that a score of 45 has been traditionally considered the cut-off for a greater risk of falls [Berg et al., 1992].

Similarly, TBI as a group showed significantly greater postural instability i.e. larger COP displacement (i.e., body sway) in response to the balance perturbation compared to HC. This balance impairment is, however, quite variable with half of our TBI population having COP values in the range of those of HC.

Despite a moderate relationship between COP and BBS (*r* = −0.52, *p* = 0.03) in our sample, the two measures may reflect different aspects of balance control. Berg et al. found that BBS was more strongly correlated with functional measures of balance such as timed up-and-go (TUG) than laboratory measures of body sway during quiet standing or in response to perturbation [Berg et al., 1992], which is consistent with our findings from our group in the same population [Pilkar et al., 2020]. Authors suggested that BBS includes various tasks that may capture several and different functional measures of balance impairment, compared to the more controlled environment of laboratory measures [Berg et al., 1992]. In comparison, the COP displacement during the balance perturbation measures the ability to dynamically correct external disturbances.

It is finally worth noting that TBI severity at time of injury doesn’t correlate and therefore is not predictive of long-term balance deficit as measured by COP displacement or BBS.

### 2. Effect of TBI on EEG brain connectivity and graph measures

Postural control is a complex task that relies on the proper functioning of distributed networks of sensorimotor-related brain regions [Takakusaki, 2017]. As TBI is often manifested as a diffuse axonal injury, the damage of the WM system can disrupt the whole-brain network connectivity and functioning [Caeyenberghs et al., 2012]. Graph theory can help identify alterations in the global brain connectivity organization [Griffa et al., 2013]. Previous studies have shown that TBI disrupts the optimal ‘small-world’ architecture of the network (as often observed in the healthy population), affecting the optimal balance between local segregation within and global integration between specialized independent subnetworks[Griffa et al., 2013]; [Zhou, 2017]; [Imms et al., 2019]. We seek to evaluate how TBI impairs this optimal integration/segregation balance at baseline and during balance perturbation.

### 3. Functional connectivity during standing

Our graph-theoretical analysis revealed significantly reduced overall connectivity strength during baseline in the theta-band but not in the alpha or beta-band in the TBI group as compared to HC. This decreased connectivity strength in lower frequency bands is in agreement with [Boshra et al., 2020], who found lower delta and theta connectivity in chronic mild TBI. Another study[Cao and Slobounov, 2010] also observed decreased long-distance connectivity from frontal areas to other parts of the brain, in the alpha band (the only explored band in the study). However, this finding was for acute mild TBI 7-days after the concussion, which may explain the discrepancies with our results. One of the common findings reported in brain disorders is the hyperconnectivity (increased functional connectivity) between certain areas within the disrupted network. However, our results contradict the hyperconnectivity hypothesis in TBI [Caeyenberghs et al., 2017] where an increase in connectivity strength or degree has been consistently found in resting-state but also task-based fMRI literature such as in [Caeyenberghs et al., 2012]; [Diez et al., 2017]. Hyperconnectivity is thought to result from an increase in local connection as a need to use detour paths around neurological disruption [Hillary and Grafman, 2017] or a compensatory response for the loss of long-range connections secondary to long-distance fiber tract damage [Venkatesan and Hillary, 2019]. The inclusion of more severe and chronic TBI in this study may explain the difference, as it has been argued that more significant disruption from severe injuries or long-term disease progression or age-related degeneration may result in hypoconnectivity (decreased connectivity) due to structural resource loss [Hillary and Grafman, 2017]; [Boshra et al., 2020]. This was illustrated in [Boshra et al., 2020] where EEG based hyperconnectivity was observed in acute mild TBI and hypoconnectivity in a chronic mild TBI. Alternatively, the brain imaging modality (e.g., EEG vs. fMRI) might be a factor confounding this observation as suggested in [Stam, 2014], especially given that TBI may alter the relationship between neural activity (EEG) and hemodynamics (fMRI) [Medaglia, 2017]. Another possible reason for the discrepancy is that the brain networks are still involved in active control of balance during quiet standing. Also, during this *baseline* period, sensorimotor networks are considered to be involved in anticipatory processing of the forthcoming postural perturbation [Varghese et al., 2019].

### 4. Task effect

When looking at the task effect on *brain activity*, we observed an overall increase during the balance perturbation in both groups across frequency bands (Fig. 5). Interestingly, the activity changes are observed in distinct brain regions supporting the view of distinct physiological roles for each frequency band. Alpha band modulation is seen over midline central (supplementary motor, lower-limb primary motor and sensory) regions, beta band over occipital and posterior parietal areas (visuo-sensory processing), and theta band over prefrontal, central and parietal areas. The role of these regions in the postural control is well-documented in the literature. For e.g., the cingulate gyrus is shown to calibrate the postural response to a challenging continuous task [Goel et al., 2019] and the role of the paracentral lobule in balance control is highlighted in [Diez et al., 2017]; [Goossens et al., 2019]; [Hagmann et al., 2008]; [Boisgontier et al., 2017], along with the precentral gyrus (M1 region) [Taubert et al., 2016]. The involvement of the superior parietal region in detecting postural instability is reported in [Slobounov et al., 2006].

More importantly, even though the band-specific spatial pattern of activation is overall consistent between groups, its amplitude is differently modulated by the task. While the increase in activity (from baseline-to-task) was reduced in alpha and beta band in the TBI group compared to HC, we observed the opposite trend for theta (i.e., TBI group showing more prominent source activity compared to HC) which points to a different neural response to the increased task demand that might indicate increased activation of another neural population.

Comparatively, we observed an overall increase in the *network connectivity* strength (NS) from baseline to task with the largest effect observed for theta (Fig. 6) consistent with [Peterson and Ferris, 2019]. The increase in NS was similar for both groups for theta, while it was significant only in HC for the alpha band, and non-significant for the beta band. This resulted in a close to significant lower alpha and beta network strength in TBI compared to HC during the task. These overall results point to impaired modulation of functional connectivity in alpha and beta. Prior literature points to the evidence that the alpha oscillations play a role in coordinating the event-related cortical processes in such a way that they inhibit task-irrelevant functional networks but facilitates task-relevant networks [Klimesch et al., 2011]. Furthermore, the function of alpha rhythms has been implicated in anticipatory sensorimotor events [Talalay et al., 2018]; [Babiloni et al., 2014]. Similarly, centro-parietal beta connectivity and activation have been implicated in attention and visuo-sensory processing to assist motor planning and execution [Chung et al., 2017]. This result is consistent with our findings regarding increased connectivity and areas of activation during the task.

Following our findings discussed above, theta-band activity has shown to increase in the frontocentral and centro-parietal regions significantly, mainly when there is an increasing balance task demand [Mierau et al., 2017]; [Solis-Escalante et al., 2019]; [Peterson and Ferris, 2018]. Fronto-central and frontoparietal theta band connections also play a vital role in postural control by increasing attention and sensorimotor processing to improve error detection and processing for the proper planning and execution of postural movements [Peterson and Ferris, 2019].

Greater activation for TBI, especially in the prefrontal area as seen in the aging population with the increasing postural challenge [Huang et al., 2017], would suggest an increased attentional demand and/or more significant cortical effort to maintain balance. We speculate this could be due to more considerable postural instability, possibly as a compensatory response to reduced connectivity. Similarly, the impaired alpha and beta neural activation and network connectivity strength modulation, without a significant impact on the balance control performance in terms of body sway, would suggest that the TBI participants were able to find a compensatory way to maintain balance for our specific balance task.

### 5. Functional Integration

Our findings showed no significant between-group differences at *baseline* in terms of the global network measures of integration, suggesting that the brain dynamics might not vary much due to pathology when there is no task demand. We observed a significant increase in theta band (but not alpha- or beta-bands) global efficiency during the task. This result is consistent with the findings in [Huang et al., 2016], where an increase in global efficiency, i.e., functional integration, was observed when participants were challenged to stand on a tilted as compared to a level-surfaced stabilometry platform. This functional integration is achieved likely through long *commissural* and a*ssociation fibers* linking different brain regions across- and within the hemisphere, respectively [Stam, 2014]. In particular, the regions that are activated along the parasagittal line of the brain (e.g., cingulate gyrus, paracentral lobule, parietal cortex) are part of a highly structurally connected backbone and can serve as a ‘hub’ of communication between other regions to facilitate functional integration [Hagmann et al., 2008]. The important integrative role of theta is consistent with the view that the neural synchronization at low frequency facilitates distant regions’ coordination, especially in response to increased task demands [Babiloni et al., 2017]. Interestingly, at baseline, the theta-band global efficiency was lower (marginally significant) in TBI than in HC, but not during the task. We could speculate that the greater increase in theta activity in TBI may reflect the greater effort needed to increase network strength and global efficiency during the task, which was lower than HC at baseline.

### 6. Functional Segregation

Concerning the functional segregation, we observed a decrease in modularity during the perturbation task compared to baseline in theta for both groups, in alpha for HC only and none for beta. More specifically, we noticed a significant reduction in the alpha modularity from baseline to task in HC but not TBI. Our conjecture is that greater cognitive demand during the task are likely to result in decreased functional segregation in healthy controls, a finding that is corroborated by[Cohen and D’Esposito, 2016]. In HC, the decrease in modularity during balance perturbation could be attributed to the increased complexity that necessitates the merging of network modules segregated during the resting state [Hearne et al., 2017]. Neuroimaging of stabilometry studies substantiate our findings [Varghese et al., 2019]; [Huang et al., 2016]. Varghese et al. [Varghese et al., 2019] postulated that the neighboring cortical areas of fronto-centro-parietal areas form new short-range connections to meet the increased demand and integration required to maintain balance control. Their findings of decreased modularity during the perturbation-related task compared to baseline were attributed to cognitive processes possibly involved in anticipatory proprioception in healthy individuals. Interestingly, there is a lack of significant change in alpha modularity in TBI compared to HC during the task. This could be the result of axonal damage in TBI, disrupting task-specific connections which might lead to difficulties in integrating different modules.

Regarding the temporal dynamics of brain activity, we construe that these networks are functionally segregated or sparsely connected during the baseline; however, they communicate with each other depending on the demand for tasks. Balance control is a complex task requiring the proper coordination of a wide visuosensory-motor network [Takakusaki, 2017]. The visuomotor and sensorimotor networks must quickly integrate the proprioceptive information from the lower limb (perturbation) in the current context of postural response to an external perturbation. As a responsive action, the supplementary motor area and associated systems try to initiate the upright posture control.

### 7. Association between the EEG graph measures and the functional outcomes (COP & BBS)

Only theta modularity was found to be negatively correlated with BBS, and this correlation was statistically significant after corrections for multiple comparisons between graph and functional measures. This suggests that a decrease in segregation would be associated with increased functional performance (i.e., higher BBS). This is consistent with the proposed important role of theta band as the conduit and SMA as a major hub of information exchange (i.e. decreased segregation) between a wide network of cognitive and visuo-sensorimotor regions to facilitate the proper detection and planning of corrective motor response to balance perturbation [Peterson and Ferris, 2019]. Similarly, in a resting-state functional connectivity study conducted by Scala et al. [Scala et al., 2019], global efficiency was negatively correlated with COP displacement suggesting that the increase in global efficiency would reflect better postural control. Theta-band modularity at rest was also recently found to be a predictor for motor learning in a sensorimotor learning task [Miraglia et al., 2018].

Our correlation results remain exploratory and caution should be exercised in interpreting this observation as causal.

### 8. Effect of TBI on structural damage

As previously mentioned, TBI can damage the WM structure due to diffuse axonal injury (DAI), potentially disrupting global functional network organization and, consequently, motor response. WM structural changes in TBI patients and healthy control were quantified using global DTI measures. Consistent with previous research studies [Douglas et al., 2015]; [Hashim et al., 2017]; [O’Phelan et al., 2018]; [Voelbel et al., 2012], we found a significant reduction in global WM microstructural integrity in TBI as reflected by lower FA and MA, and higher MD compared to controls. Reduced WM integrity and tract degeneration have consistently been reported in chronic TBI [Caeyenberghs et al., 2014]; [Hashim et al., 2017]; [Wallace et al., 2018]. Following DAI, WM tract degeneration would lead to greater free water molecules movements, resulting in lower FA and higher MD. Furthermore, reduced MA would signify more disc-like, less cigar-like shape water diffusion, i.e., more fiber crossing and less unidirectional fiber bundle [Douaud et al., 2011]; [Yoncheva et al., 2016], which could reflect the deterioration and thinning of WM tracts.

Furthermore, using TBSS, we compared the WM tracts in skeletonized maps in TBI and HC as a voxel-based group-level analysis for all three measures. Although global FA, MD, and MA were significantly changed in TBI, when these measures were registered on the standard space of major fiber tracts and projected onto the mean FA skeleton, significant group differences were found only in FA, but not in MA or MD. In FA skeleton maps, the main regions affected were the thalamus and some parts of the Corpus Callosum, and parietal-occipital junction regions, which is consistent with prior findings [O’Phelan et al., 2018]; [Veeramuthu et al., 2015]; [Hashim et al., 2017]; [Owens et al., 2018]. These findings have important applications given the role of the corpus callosum in cross-hemisphere communication and error detection in balance perturbation [Peterson and Ferris, 2019]; [Peterson and Ferris, 2018] and thalamus in conveying all types of sensory information to the cortex to perform balance and other sensory-motor functions [Surgent et al., 2019]. Future analyses should further investigate the regional and tract-based WM damage and their potential role in balance deficit and abnormal postural response to the perturbation task.

### 9. Association between DTI and BBS, and DTI and graph measures

Correlation analysis between the DTI measures and EEG graph-theoretic measures revealed that there was a consistent relation between the structural damage and less integrated, global synchronous connectivity during the task in TBI in alpha and beta bands. Particularly, the correlation between the alpha-band network strength and the white matter structural integrity captured by the global FA was close to the conservative Bonferroni-corrected significance level. The long-range connectivity was shown to be associated with the fractional anisotropy, and the damage to these long-range connections was reflected in the global efficiency values [Rudie et al., 2013].

Moreover, the negative correlation between the Berg Balance Scale and mean diffusivity hints at overall WM degeneration in poorer balance performance. Given our finding is based on the global mean diffusivity, our future work will investigate the diffusivity characteristics in more local tracts such as middle and inferior cerebellar peduncles, shown to be involved in postural control [Caeyenberghs et al., 2010].

## Limitations and Future directions

We acknowledge that our study has some limitations, particularly with the small sample size. Also, there is another limitation concerning the heterogeneity within the sample population; this is particularly challenging in the TBI group because of the varying degrees of injury (mild/moderate/severe) not necessarily correlating with the motor deficits. However, we have tried our best to control for other factors such as age, height, and weight, which contribute to the postural control performance.

We cannot completely rule out the potential effect of motion artifacts on our results especially for low frequencies [Nordin et al., 2019]. However, source-localized measures and phase coherence rather than channel-based amplitude-based connectivity or activity, are more immune to motion artifacts. Also, the band-specific rather than a broad spatial pattern of activity modulation (Fig. 5) as well as the negative rather than positive correlation between COP (movement) and integration, would suggest a true neural response to the task.

While we integrated information available from both task-based neuroimaging using the EEG and the structural imaging using DTI, it is worth highlighting that the functional connectivity between the time-series of two regions is often modulated by factors independent from the structural connectivity. A functional connectivity measure between two areas can be influenced by the third region with no direct fiber track in between, but still coordinated functionally via cortico-thalamocortical pathways [O’Reilly et al., 2017]. We recommend that future studies investigate the causal association between structural and functional connections.

We admit we did not compare the graph metrics to the null model in our graph-theoretic analysis. In our defense, we highlight that the choice of the null model plays a critical role, and it can dramatically change the small-world properties of the network [Fallani et al., 2014]. Moreover, the choice of FC measurement (e.g., correlation/coherence) can also affect the comparison concerning the null model. Nevertheless, unlike the correlation-based FC measurement, which often results in a spurious association between two regions, the coherence-based FC is tested for its statistical significance using Schelter’s approach [Schelter et al., 2006]. Incorporating these methodological details is our priority in the upcoming graph-theoretic studies.

In this study, our focus has been primarily on the global measures of EEG connectivity graphs. However, the underlying neural mechanisms of the postural control are too complex to summarize with a single statistical value. Depending on the research question, this can be either a strength or a weakness. In our current approach, we attempted to address whether we can identify a type of neural biomarker based on the EEG connectivity graph. While the global graph measures allow us to explore the association between the connectivity-based graph measures and the functional outcomes, it still does not answer questions related to the local characteristics of task-relevant networks. Notwithstanding this limitation, we did explore the local properties based on the weighted node strength and the significant cortical activity in the regions of interest to the postural control. Future work will examine the significance of these nodes in terms of local graph properties such as rich-club coefficient and centrality measures.

## Conclusions

To the best of our knowledge, this is the first study investigating the brain network segregation and integration in a TBI population during a postural control task performed on a computerized stabilometric platform. The findings from EEG graph measures in TBI compared to healthy controls revealed altered baseline and task modulation of global graph-theoretic measures of *network strength, global efficiency,* and *modularity* in the brain functional networks. Interestingly, reduced network connectivity strength and integration, and greater network segregation were correlated with poorer balance performance (COP and BBS) and greater structural brain damage. Findings from the graph measures were found to be frequency bands and region-specific, thus highlighting their distinct role.

The combined use of EEG-based FC measures during the task and the DTI-based structural integrity measures helped provide new insights into the underlying structural-functional mechanisms of postural control in TBI. These observations could pave the way for future research to identify cortical biomarkers of postural control deficits in TBI, thus potentially assisting clinicians and researchers to better understand neuromuscular disorders.

We believe that our findings and inferences from this pilot study should provide directions to future studies on brain connectivity in TBI and other neuropathologies.

## Data Availability

All the raw data are not publicly available due to the IRB restrictions. However, the MATLAB scripts and the de-identified pre-processed EEG data supporting this study can be made available upon reasonable requests to the corresponding author.

