## Supplementary Material for "Graph-theoretical Analysis of EEG Functional Connectivity during Balance Perturbation in Traumatic Brain Injury: A Pilot Study"

### S1. EEG Source localization:

EEG Source Localization (ESL) allows one to study the cortical dynamics using the underlying cortical sources estimated from the sensor-space EEG (Michel & Brunet, 2019; Handiru et al., 2018). The ESL problem is mathematically formulated as:

$$\mathbf{X} = \mathbf{L}_f \mathbf{S} + \epsilon \tag{1}$$

where $\mathbf{X} \in \mathbb{R}^{N \times T}$ represents the sensor-space EEG with $N$ channels (or sensors) and $T$ timepoints, $\mathbf{L}_f \in \mathbb{R}^{N \times M}$ denotes the lead-field matrix that projects $M$ sources $(M \gg N)$ onto the sensor-space, $\mathbf{S} \in \mathbb{R}^{M \times T}$ denotes the cortical source time series, and $\epsilon \in \mathbb{R}^{N \times T}$ denotes the measurement noise. The two unknowns, $\mathbf{L}_f$ and $\mathbf{S}$ are estimated using (1) forward modeling and (2) inverse modeling, respectively. Forward modeling is done to compute the volume conductor model which realistically approximates the electromagnetic field propagation through different layers in the head model (scalp, skull, cerebrospinal fluid, grey matter, and WM). As the sensor-space EEG is only a projection of the true cortical activity, one needs to compute a realistic individual head model for which we use the subject-specific anatomical data. To this end, we used a subject-specific T1-weighted MRI and 3D digitized EEG positions to compute the head model by means of the 3-layered Boundary Element Method (BEM) (Fuchs et al., 1998). The default settings of conductivity values in OpenMEEG software were used for 3-layered BEM (scalp =1, skull = 0.0125, brain = 1) (Gramfort et al., 2010). Once we obtained the head model, $\mathbf{S}$ was computed using an inverse modeling algorithm named sLORETA (standardized Low-Resolution Electromagnetic Tomography) (Pascual-Marqui, 2002). All the aforementioned steps were implemented using Brainstorm software (Tadel et al., 2011).

### S2. Functional connectivity:

Functional connectivity was computed using the imagery part of coherence as a measure of phase synchrony between two brain signals. We denote the imaginary part of coherence as $\text{Im}(\underline{\mathbf{\Gamma}})$ with $\underline{\mathbf{\Gamma}} \in \mathbb{C}^{U \times U \times V}$ being a three-way tensor, where $U$ and $V$ are the number of ROIs and frequency bins of interest, respectively. The phase synchrony between two signals $i$ and $j$ can be measured using $\mathrm{Im}(\underline{\boldsymbol{\Gamma}}(i, j, :))$, which has been employed in Nolte 2004 to represent the statistical similarity between two brain sources derived from EEG. It is desirable to use $\mathrm{Im}(\underline{\boldsymbol{\Gamma}})$ since one can easily isolate the entries arising due to the volume conduction effect (VCE), i.e., the non-zero values of the tensor guarantee non-zero delays. In other words, the entries of $\mathrm{Im}(\underline{\boldsymbol{\Gamma}}(i, j, :)) \neq 0$, only when the respective frequency components of $i$ and $j$ are statistically independent. Therefore, the non-zero entries of $\mathrm{Im}(\underline{\boldsymbol{\Gamma}}(i, j, :))$ imply that there is no time-lagged interaction between the corresponding frequency bins of $i$ and $j$ due to VCE. Note that $i$ and $j$ are the time series generated by the brain ROIs of interest in our context.

Let us consider the $k^{\mathrm{th}}$ segment of the $i^{\mathrm{th}}$ time series as $x_{i,k}$ and its corresponding Fourier Transform as $X_{i,k}(f)$ at frequency $f$. The cross-spectral matrix $S_{i,j}(f)$ is then computed as (Nolte et al., 2004).

$$S_{i,j}(f) = \frac{1}{K} \sum_{k=1}^{K} X_{i,k}^{*}(f)\, X_{j,k}(f) \tag{2}$$

$$\underline{\boldsymbol{\Gamma}}(i, j, f) = \frac{S_{i,j}(f)}{\left[S_{i,i}(f)\, S_{j,j}(f)\right]^{1/2}} \tag{3}$$

where * refers to the complex conjugate.

### S3. Beta Band Results:

Table 1: Summary of 2-way repeated-measures ANOVA of Beta-band graph measures

| | Group | | | Time | | | Group x Time | | |
|---|---|---|---|---|---|---|---|---|---|
| | F(1,30) | p | Effect Size | F(1,30) | p | Effect Size | F(1,30) | p | Effect Size |
| Network Strength | 4.67 | 0.039 | 0.135 | 0.69 | 0.41 | 0.023 | 0.03 | 0.86 | 0.001 |
| Global Efficiency | 1.039 | 0.32 | 0.033 | 0.567 | 0.46 | 0.019 | 0.62 | 0.44 | 0.02 |
| Modularity | 3.00E-04 | 0.98 | 9.00E-05 | 2.67 | 0.11 | 0.082 | 0.056 | 0.81 | 0.002 |

Effect size is denoted using partial eta squared values ($\eta_p^2$)

Table 2: Descriptive statistics and contrast analysis of beta-band graph measures

| | | | | |
|---|---|---|---|---|
| Network Strength | TBI | 461 ± 39 | 466 ± 29 | t=0.49 , p=0.63, d=0.1 |
| | HC | 474 ± 22 | 484 ± 36 | t=0.83, p=0.42, d=0.21 |
| | Between Group | t=1.23, p=0.22, d=0.44 | t=1.51, p=0.15, d=0.53 | |
| Global Efficiency | TBI | 0.478 ± 0.025 | 0.49 ± 0.036 | t=1.34, p=0.32 , d=0.23 |
| | HC | 0.476 ± 0.025 | 0.476 ± 0.026 | t=5.62, p=0.98, d=0.007 |
| | Between Group | t=0.23, p=0.81, d=0.08 | t=1.16, p=0.26, d=0.41 | |
| Modularity | TBI | 0.02 ± 0.002 | 0.017 ± 0.001 | t=1.26, p=0.21, d=0.36 |
| | HC | 0.019 ± 0.002 | 0.018 ± 0.001 | t=0.7, p=0.48 , d=0.23 |
| | Between Group | t=0.13, p=0.89, d=0.35 | t=0.15, p=0.88, d=0.05 | |

Between Group (df=30); Within-group TBI (df=32), Within-group HC (df=28), d: Effect size is denoted using cohen's D value.

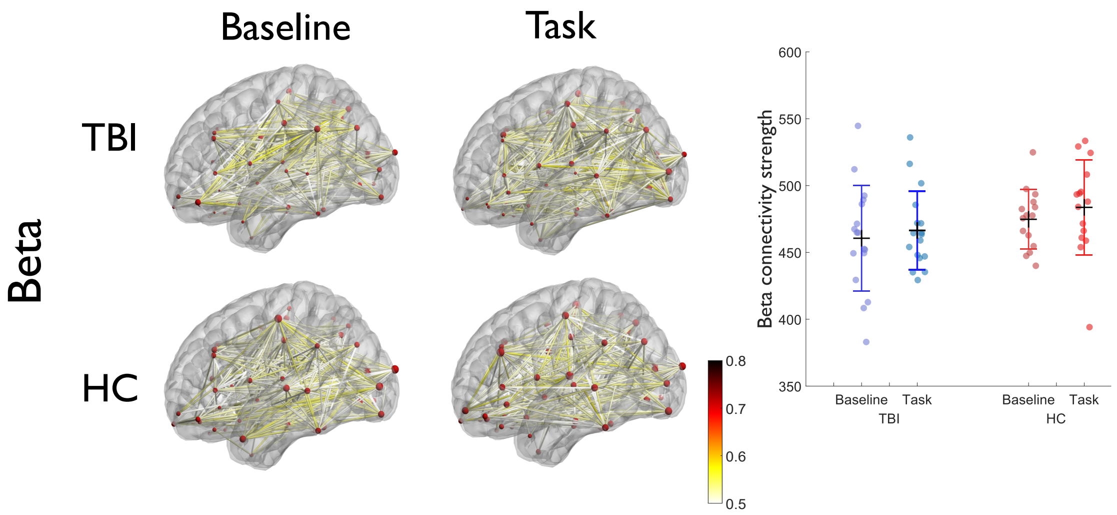


Figure 1: Whole-brain functional connectivity based on source-space EEG coherence in beta-band in each group (*TBI* and *HC*) across time periods (*baseline* and *task*). Group-level connections are averaged and plotted as an edge between different ROIs anatomically parcellated using Desikan-Killiany Atlas. For better visualization, the connections are thresholded at edge weight = 0.3. Seed voxels of each ROI are indicated as spheres with a radius proportional to their node strength. On the right side, the boxplot comparison of network strengths during the baseline vs. task period is shown for both the groups. No statistically significant differences were found in within-group as well as between-group comparisons.

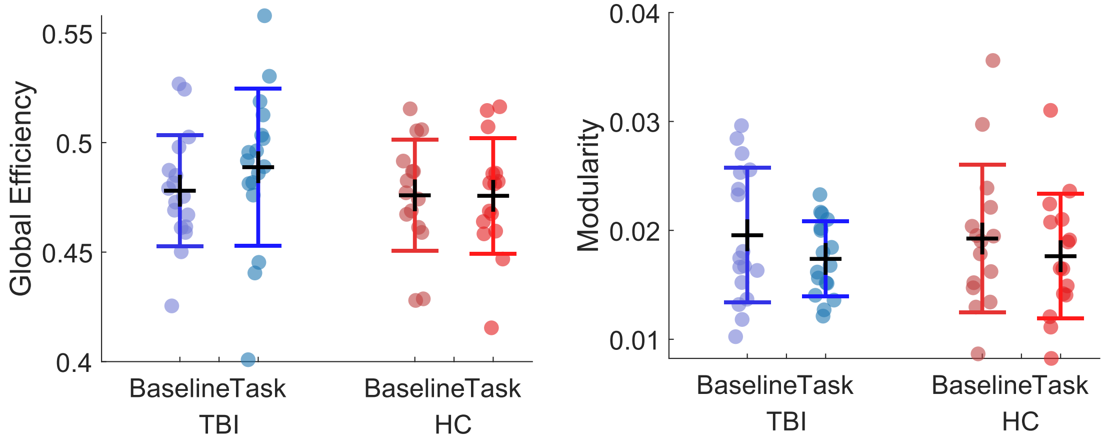


Figure 2: Group-level comparison of beta-band network measures of functional integration (left) and functional segregation (right). The categorical scatter plot of the network measures during the baseline and during perturbation (task) within each group is shown above. The black horizontal line indicates the mean and the colored horizontal lines indicate the standard deviation. No statistically significant results were found in any of the comparisons.

#### Association between DTI and Graph Measures:

The global FA was negatively correlated with the alpha-band network modularity ($r$ = -0.65, $p$ = 0.03 without and $r$ = -0.59, $p$ = 0.04 with outliers), and positively correlated with the GE ($r$ = 0.59, $p$ = 0.04)(Fig. **??**b). Although the beta-band network measures did not show any significant effects earlier in terms of the within-group and between-group differences, we observed the beta-band network strength was significantly associated with all the DTI measures in the TBI group (Fig. **??**c). The beta-band network strength was positively associated with the global FA ($r$ = 0.72, $p$ = 0.01 without and $r$ = 0.65, $p$ = 0.02 with outliers) , and negatively correlated with the global MD ($r$ = 0.71, $p$ = 0.01 without and $r$ = -0.64, $p$ = 0.047 with outliers).
